# Investigation of the impact of bromodomain inhibition on cytoskeleton stability and contraction

**DOI:** 10.1101/2023.11.14.567076

**Authors:** Alexander Bigger-Allen, Ali Hashemi Gheinani, Rosalyn M. Adam

## Abstract

Injury to contractile organs such as the heart, vasculature, urinary bladder and gut can stimulate a pathological response that results in loss of normal contractility. PDGF and TGFβ are among the most well studied initiators of the injury response and have been shown to induce aberrant contraction in mechanically active cells of hollow organs including smooth muscle cells (SMC) and fibroblasts. However the mechanisms driving contractile alterations downstream of PDGF and TGFβ in SMC and fibroblasts are incompletely understood, limiting therapeutic interventions. To identify potential molecular targets, we have leveraged the analysis of publicly available data, comparing transcriptomic changes in mechanically active cells stimulated with PDGF and TGFβ and identified a shared molecular profile regulated by MYC and members of the AP-1 transcription factor complex. We also analyzed data sets from SMC and fibroblasts treated in the presence or absence of the MYC inhibitor JQ1. This analysis revealed a unique set of cytoskeleton-associated genes that were sensitive to MYC inhibition. JQ1 was also able to attenuate TGFβ and PDGF induced changes to the cytoskeleton and contraction of smooth muscle cells and fibroblasts *in vitro*. These findings identify MYC as a key driver of aberrant cytoskeletal and contractile changes in fibroblasts and SMC, and suggest that JQ1 could be used to restore normal contractile function in hollow organs.

## Introduction

The function of hollow organs such as the heart, gut, vasculature and urinary bladder rely on coordinated cycles of contraction and relaxation. In response to injury, the normal contractile activity of mechanically active cells such as smooth muscle cells, fibroblasts and myofibroblasts, is perturbed leading to aberrant contraction and functional decline [1]. In addition to actomyosin cross-bridge cycling, tension generation in cells involves contributions from the actin cytoskeleton, as evidenced by the reduction in smooth muscle contraction following pharmacological inhibition of actin polymerization (reviewed in [2]). In spite of extensive study of mechanisms that regulate the actin cytoskeleton, exploiting this knowledge to reverse aberrant contraction remains challenging.

Previous studies from our group identified the AP-1 transcriptional complex as a key regulator of smooth muscle cell behavior in response to discrete stimuli such as cyclic stretch-relaxation and the physiologically relevant growth factors Transforming Growth Factor beta (TGFβ) and Platelet Derived Growth Factor (PDGF) [3-8]. TGFβ, a known inducer of smooth muscle differentiation and activator of fibroblast to myofibroblast transdifferentiation [9, 10] was found to induce the formation of filamentous actin in smooth muscle cells [7]. In that study, knockdown of the AP-1 monomer JUNB decreased both basal and TGFβ-stimulated cell contraction and cytoskeletal tension, in parallel with reduction in phosphorylation of both Myosin Light Chain 20 (MLC20) and cofilin (CFL). Of note, these changes occurred independently of alterations in the SM markers; alpha Smooth Muscle Actin (αSMA), Smooth Muscle protein 22 (SM22α) and Calponin 1 (CNN1). We also implicated Activator Protein 1 (AP-1) in regulation of gene expression and Smooth muscle cell (SMC) phenotype downstream of Platelet Derived Growth Factor Receptor (PDGFR) -mediated signaling [5, 8]. In that study an unbiased assessment of PDGF-induced transcriptomic changes in SMC identified AP-1 and MYC proto-oncogene (MYC) as the transcriptional regulators most highly linked to up-regulated genes [8]. In addition, integration of gene expression data with proteomics led to the identification of a novel MYC-centric network in SMC. Moreover, manipulation of the MYC target and RHOA effector DIAPH3, by siRNA or with a MYC inhibitor, led to marked cytoskeletal changes in SMC consistent with prior reports linking DIAPH3/mDia2 to RHOA-dependent regulation of SMC phenotype [11, 12].

Previous studies have explored the functional relationship between MYC and actin cytoskeleton regulation in non-muscle cells [13-18]. Whereas some reports have identified MYC as an inhibitor of actin filament polymerization [15], others have demonstrated enhanced formation of F-actin stress fibers [18]. However, since these analyses were conducted under conditions where MYC was overexpressed, the significance of the functional relationship between MYC and cytoskeletal regulation in non-transformed cells with endogenous MYC expression was unclear. To address this gap in knowledge, a number of groups have explored the impact of pharmacological inhibition of endogenous MYC. Among the inhibitors assessed, JQ1 has been shown to attenuate MYC expression indirectly by preventing the bromo and extra-terminal (BET) family of proteins from binding to acetylated histones [19]. Although studied primarily in the context of MYC overexpression in cancer, JQ1 and other BET protein inhibitors have also been employed in non-cancer settings including fibrosis and cardiac diseases. In these studies JQ-1 treatment attenuated epithelial-to-mesenchymal transition, migration and activation of myofibroblasts consistent with an impact on the cytoskeleton [20]. However, the extent to which inhibiting MYC may be beneficial in settings of aberrant contraction remains unclear. In this study, we have identified MYC as a central mediator of pathophysiologically relevant stimuli in smooth muscle cells and fibroblasts. Additionally, we have explored the impact of MYC inhibition on the regulation of the actin cytoskeleton and contraction in these mechanically active cells.

## Results

### In silico analysis reveals common transcriptional regulators in mechanically active cells

Prior findings from our groups implicated PDGF and TGFβ as drivers of key pathways in SMC relevant to pathological tissue remodeling [5, 6, 7, 8, 21, 22]. MYC and AP-1 emerged as functionally relevant mediators, although the extent to which they were regulated in common by PDGF and TGFβ remains unknown. To investigate effectors shared between PDGF-or TGFβ- treated bladder SMC, we re-analyzed gene expression data generated previously by us [8], and compared the results with re-analyzed publicly available data of SMC and fibroblasts stimulated with PDGF and TGFβ. After performing filtering assessments of the publicly available data **(Supplemental Figure 1),** we identified nearly 800 differentially expressed genes (DEGs) in common between our and these three other datasets. **(Figure 1A, D, and G)**. To better understand the regulatory network evoked by PDGF and TGFβ treatment, we used the ChEA3 transcription factor (TF) enrichment analysis tool [23] to identify TFs that could explain the gene expression profiles. This analysis identified MYC and the MYC-associated factor X (MAX) among the most enriched TFs, regulating the three gene sets of roughly 800 genes from each comparison **(Figure 1B, E, and H**). Additional highly-enriched TFs included members of the AP- 1 complex such as FOS, FOSB, JUN and/or JUND, as well as E2F4 and NFYA.

**Figure 1:**
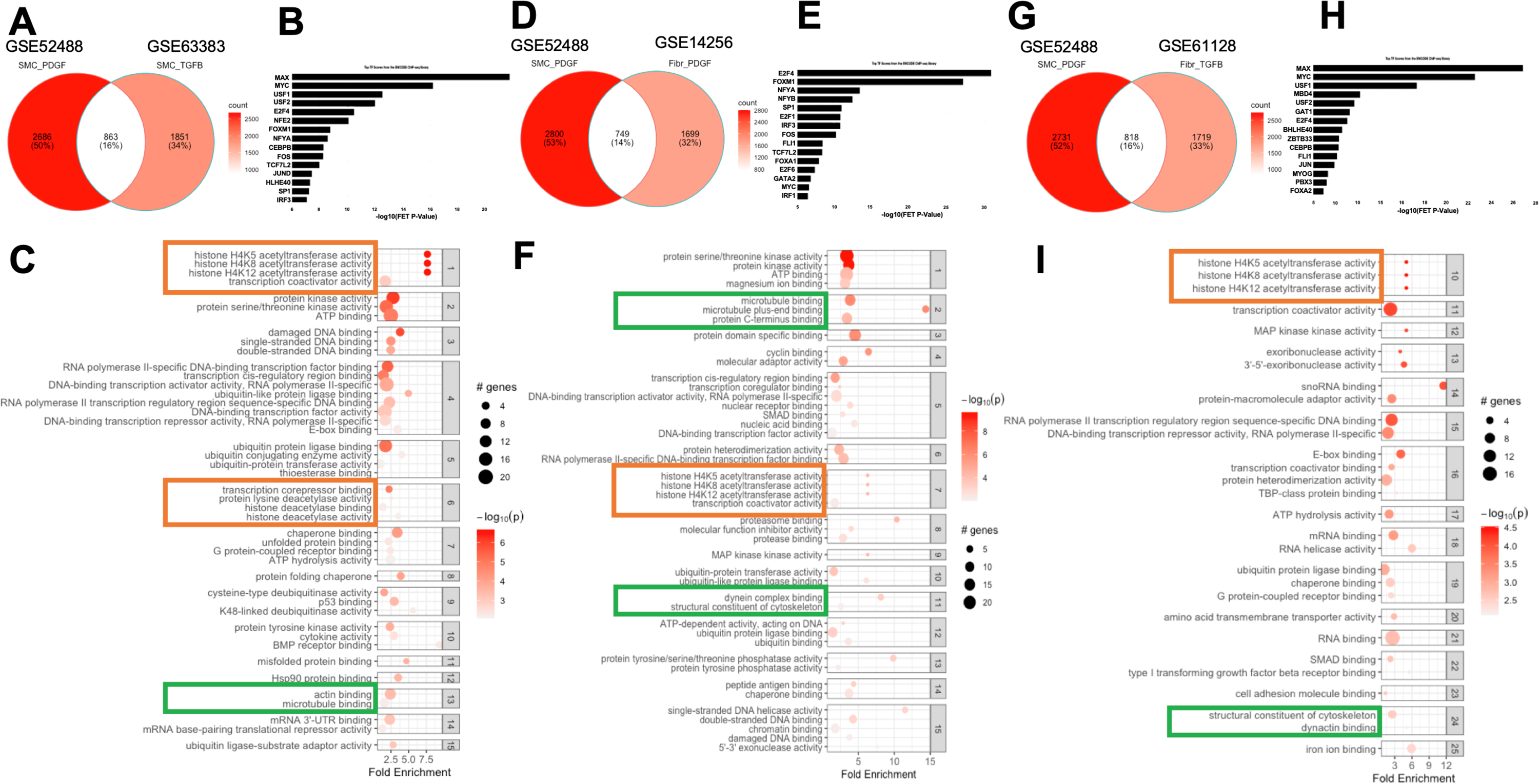
Comparative analysis of growth factor stimulated mechanically active cells identifies common TF regulators and changes in chromatin and the cytoskeleton **(A, D, G)** Venn diagrams comparing our PDGF stimulated pHBSMC DEGs to the DEGs generated from reanalyzed publically available data of smooth muscle cells or fibroblasts stimulated with PDGF or TGFB. **(B, E, H)** Transcription factor master regulator analysis using CHEA3 and ENCODE database of all the shared DEGs from A, D, and G identified MYC, MAX and members of the AP-1 transcription factor complex as the most highly networked regulators based on -log Fisher’s exact test (FET) p-value. **(C, F, I)** Shared DEGs from A, D, and G were subjected to enrichment analysis using GO terms for molecular function. Terms were then clustered to group molecular functions with overlapping DEGs. Highlighted in orange, are molecular functions related to chromatin remodeling. Highlighted in green are molecular functions related to cytoskeleton regulation.

To understand what processes these genes are involved in, we performed enrichment analysis on the shared DEGs from each of the three comparisons utilizing gene ontology (GO) terms for molecular function (MF) **(Figure 1C, F and I),** cellular compartment (CC) **(Supplemental Figure 2A, B, and C)**, and biological process (BP, data not shown). In the first comparison, our previously generated data were compared to SMC stimulated with TGFβ. The top terms enriched downstream of shared DEGs from PDGF and TGFβ treated SMC include histone acetyltransferase and deacetylase activity in clusters 1 and 6 (7.66-fold enrichment, p-value = 9.29e-05 and 1.33-3.51-fold enrichment, p-value = 0.005 – 1.27e-05 respectively) **(Figure 1C)**. Similarly, enrichment analysis of shared DEGs between our dataset and GSE14256, in which fibroblasts were stimulated with PDGF, identified histone acetyltransferase activity in cluster 7 (6.26-fold enrichment, p-value = 4.31e-05) (**Figure 1F)**. Further, histone acetyltransferase activity was also identified in cluster 10 when comparing our data with GSE61128 in which fibroblasts are stimulated with TGFβ (5.02-fold enrichment, p-value = 0.0063) **(Figure 1I)**. The enrichment of histone acetyltransferase activity in all comparisons supports a shared induction of chromatin-related genes by both PDGF and TGFβ in SMC and fibroblasts.

In all three comparisons of our data with fibroblasts and SMC stimulated with TGFβ or PDGF, the perturbation to genes associated with cytoskeleton was also notable. To understand the extent to which all four datasets shared DEGs, we generated a 4-way Venn diagram and identified 89 shared genes (**Supplemental Figure 3A).** Using the ChEA3 TF enrichment tool, MYC and MAX were predicted as the top most networked TF regulators (**Supplemental Figure 3B)**. Finally, gene set enrichment analysis was performed using terms for MF, CC and BP terms **(Supplemental Figure 3C, D, E)**. In the CC enrichment analysis, positive regulation of cytoskeleton organization was one of the most enriched terms, comprising 5 genes (LIMK1, BIN1, CDKN1B, BCAS3, and WASL, gene list not shown) shared between all 4 datasets **(Supplemental Figure 3D)**. Taken together these analyses implicate MYC and MAX as shared drivers that mediate PDGF-and TGFβ-induced changes to the cytoskeleton.

### Transcriptomic data from JQ1 treated mesenchymal cells support JQ1 attenuation of cytokine induced changes

Given the identification of MYC and MAX as highly enriched TFs downstream of both PDGF and TGFβ treatment, we next explored the impact of JQ-1 treatment on gene expression profiles from mechanically active cells exposed to proliferative or pro-contractile stimuli. JQ-1 is a bromodomain inhibitor that has been shown previously to inhibit MYC-dependent transcription [24, 25] and a reduction in MYC protein stability [26]. In addition, JQ1 has previously been demonstrated to inhibit multiple members of the AP-1 TF complex by reducing both total JUN expression and JUN phosphorylation [27-30]. We identified and re-analyzed 4 publicly available datasets in which SMC or fibroblasts were treated without or with JQ-1: rat vascular SMCs exposed to PDGF (GSE111714); human vascular SMC stimulated with undefined growth medium (GSE138323); rat cardiac fibroblasts stimulated TGFβ1 (GSE127229)[31]; and Rheumatoid Synovial Fibroblasts stimulated with IL1β (GSE148395)[32]. PCA plots of count and expression matrices of the publicly available data were visualized to determine quality of replicates and separation of conditions **(Supplemental Figure 4)**. Greater than 60% of the variation for each dataset was accounted for by the first two principal components demonstrating that most of the differences were due to experimental stimulations. DEGs from the JQ1 datasets were analyzed using the degPatterns function, which identified two groups of genes from each dataset: those that were either up-or down-regulated with the stimulus, and subsequently attenuated with JQ1 co-stimulation (**Figure 2A, C, E, and G)**. Genes that fit this pattern were classified as JQ1-sensitive genes.

**Figure 2.**
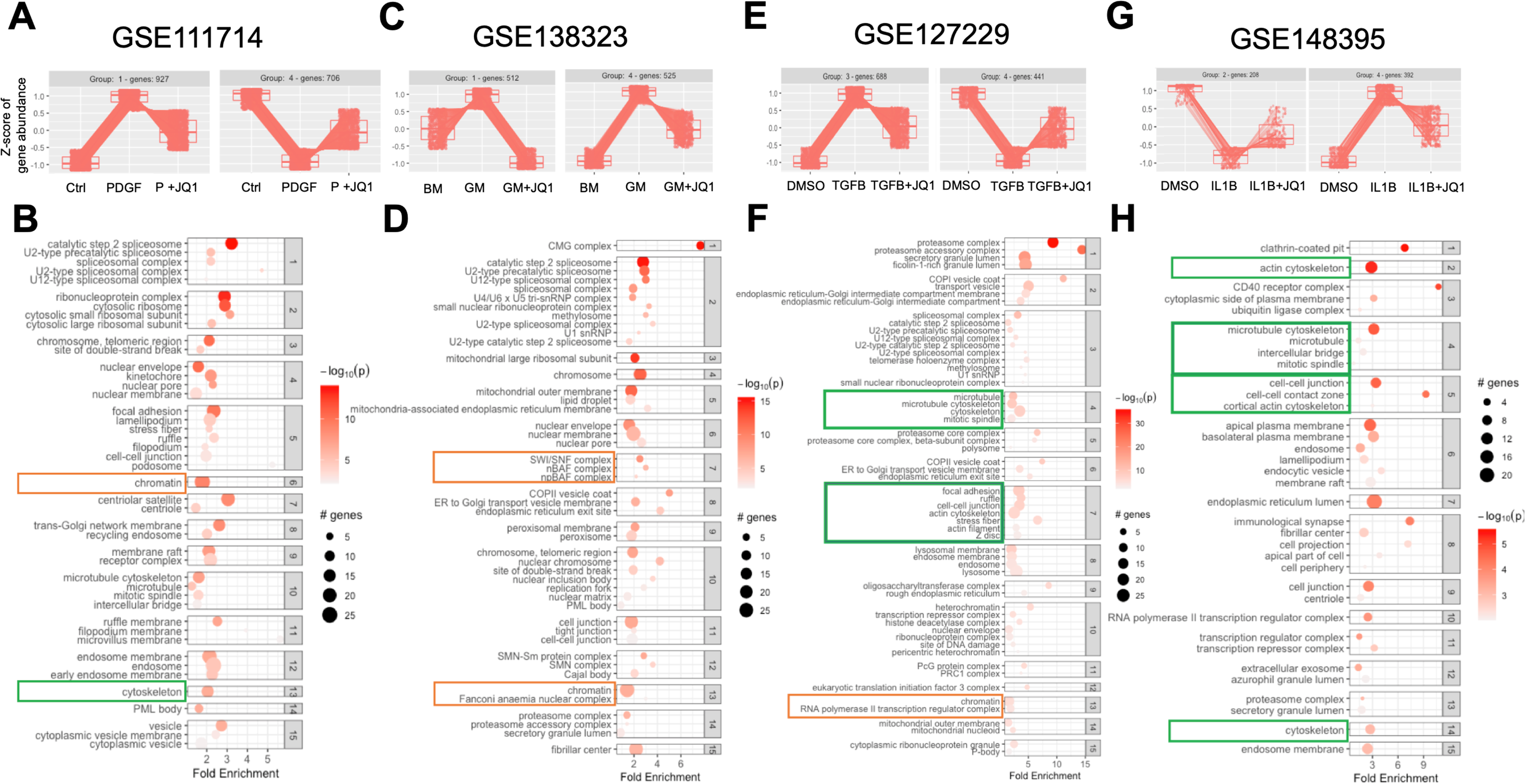
Pattern and enrichment analyses support JQ1 attenuation of genes associated with cytoskeletal changes. Four publicly available datasets in which mechanically active cells were stimulated with a physiologically relevant pro-fibrotic condition in the absence or presence of JQ1 were identified and reanalyzed. Differential gene expression analysis was performed using DESeq2 after generating count matrices from the fastq files using Biogrids software tools including: SRAtoolkit, STAR, Subread. (**A, C, E, G)** Pattern analysis of DEGs from each of the four datasets identified 2 groups of genes that were perturbed by PDGF (GSE11714), Growth medium (GSE138323), TGFB (GSE127229) or IL1B (GSE148395) and attenuated by JQ1. (**B, D, F, H)** JQ1 sensitive genes were subjected to GO cellular compartment terms and clustered based on common genes between terms. Highlighted in yellow are terms related to chromatin remodeling. Highlighted in green are terms related to cytoskeleton regulation.

Enrichment analyses on JQ-1-sensitive DEGs was performed using GO-CC terms (**Figure 2 B, D, F and H)** as well as GO-MF terms **(Supplemental Figure 5A-D**). We combined the two sets of JQ1-sensitive DEGs identified in each dataset for this analysis. In rat VSMCs treated with PDGF (GSE111714), JQ1 attenuated changes related to chromatin remodeling (cluster 6, fold enrichment = 1.78, p-value = 9.95e-10) and the cytoskeleton (cluster 13, fold enrichment = 2.04, pvalue = 7.77e-03) (**Figure 2A)**. Similarly, in human VSMC stimulated with growth media (GSE138323), JQ1 attenuated changes corresponding to chromatin remodeling (cluster 7, fold change = 2.50, pvalue = 2.30e-09 and cluster 13, fold change = 1.41, pvalue = 1.27e-07). In cluster 39 (data not shown), genes involved in the actin cytoskeleton were enriched (fold change = 1.28, pvalue = 8.91e-04) **(Figure 2B)**. Although the datasets describe two different treatments in VSMC from two species, JQ1 was able to attenuate expression of genes associated with chromatin remodeling and the actin cytoskeleton.

Transcriptomic datasets GSE127229 and GSE148395 were generated from rat cardiac fibroblasts treated with TGFβ1 +/-JQ1 and human rheumatoid synovial fibroblasts stimulated with IL1β +/- JQ1, respectively, and analyzed as previously mentioned. In both datasets two groups of JQ1 sensitive genes were identified in which TGFβ or IL1β treatment perturbed gene expression compared to vehicle (DMSO)-treated control and were attenuated by JQ1 (**Figure 2 E and G)**. These groups of JQ1 sensitive genes were combined for each dataset to perform ontology enrichment analysis for GO-CC and GO-MF terms. Both chromatin and cytoskeleton-related changes were enriched in the JQ1 sensitive genes in both fibroblast datasets. Notably, the cytoskeleton-associated processes were more highly enriched in both fibroblast datasets compared to the SMC datasets. Additionally, these cytoskeleton-associated terms were more highly enriched than chromatin-associated terms within each dataset compared. This further supports the hypothesis for a novel mechanism whereby treatment with JQ1 can regulate effectors of the cytoskeleton not only within fibroblasts, but also in SMC albeit to a lesser extent.

Next, the expression of a list of cytoskeleton-related genes was compared between the datasets to identify those that were attenuated by JQ1 compared to the cytokine treatment alone **(Supplemental figure 6A)**. We excluded the microarray dataset (GSE138323) of SMC stimulated with undefined growth media since all JQ1-sensitive genes were induced by growth media, and none were down regulated, precluding a robust comparison of genes based on direction of differential expression and attenuation. Among the 3 remaining datasets there were 8 JQ1-sensitive genes that were induced by cytokine (PDGF, TGFβ, or IL1B) and reduced by JQ1. Similarly, there were 9 genes that were reduced by PDGF or TGFβ whose decrease was attenuated by JQ1. To visualize these JQ1-sensitive cytoskeleton-associated genes we utilized the KEGG pathway Regulation of Actin Cytoskeleton and mapped the JQ1-downregulated genes in blue and the JQ1-induced genes in red **(Supplemental Figure 6B)**. These genes, listed in the tabular view, represent enzymes and structural components of the cytoskeleton **(Supplemental Figure 6C)**.

### *In vitro* validation of JQ1-sensitive cytoskeletal targets

To validate predictions from the *in silico* analysis we explored sensitivity of of a subset of cytoskeleton-associated genes to JQ1 in rat bladder mesenchymal cells (RBMC) and human bladder SMC **(Figure 3A).** Cells were stimulated with PDGF or TGFβ with or without JQ1 for 0.5, 2, 4 or 16 hrs, and expression of 6 genes predicted to be induced by PDGF and TGFβ in the publicly available datasets and sensitive to JQ-1 **(Supplemental Figure 6A)** were assessed by qRT-PCR: Platelet derived growth factor receptor alpha (Pdgfrα), Integrin beta 1 (Itgb1), Integrin alpha 5 (Itga5), actinin 1 (Actn1), Src proto-Oncogene, non-receptor tyrosine kinase (Src), Beta actin (Bact), Lim kinase (Limk) and cofilin (Cfl). Notably, PDGF and TGFβ induced expression of all 8 targets at 2 hrs (1.34, 1.43, 0.97, 1.43, 1.60, 1.01, 0.60, 1.00 and 0.63, 1.05 1.02, 1.28, 1.68, 1.13, 0.83, 0.88 log2FC respectively). (**Figure 3A**). All targets except, Bact and Acnt1 were suppressed to near baseline levels by JQ1 as early as 2 hrs post stimulation with PDGF or TGFβ. By 16 hrs, JQ1 suppressed PDGF and TGFβ induction of only Pdgfrα, Itga5, Src and Actn1. pHBSMC showed similar reduction in all target gene expression with JQ1 by 4hrs regardless of the PDGF or TGFβ stimulation **(Supplement Figure 8).** Overall, the validation of JQ1 sensitivity of a subset of cytoskeletal effectors suggests a role to use JQ1 to target MYC regulated cytoskeletal changes.

**Figure 3.**
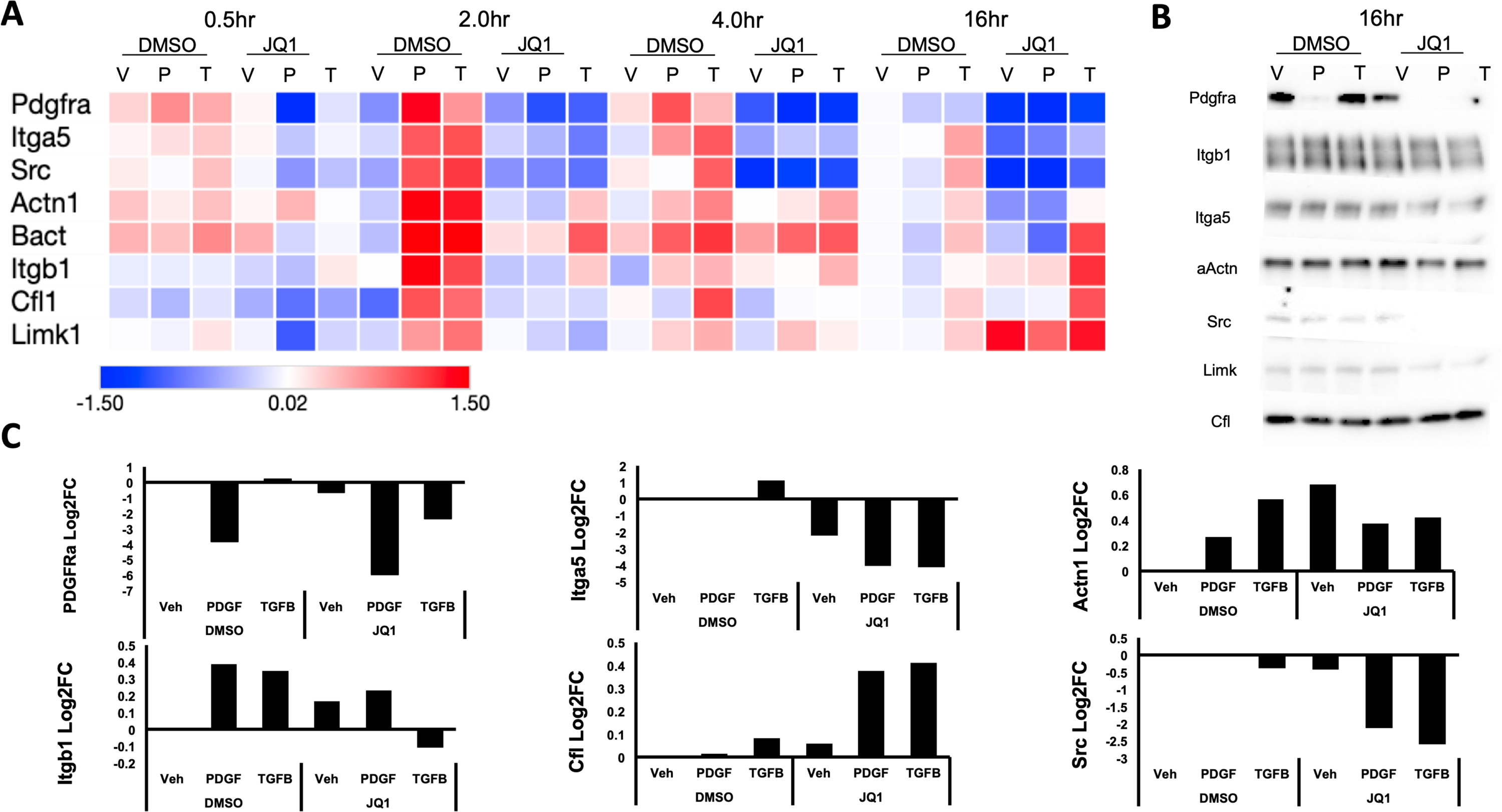
Validation of JQ1 sensitive cytoskeletal effectors via qPCR and immunoblots. A subset of cytoskeleton associated targets predicted to be sensitive to JQ1 were validated using qPCR and immunoblotting techniques. Primers for JQ1 sensitive cytoskeleton associated targets were generated using NCBI primer design tool. (**A)** A time course of PDGF and JQ1 concurrent stimulation of RBMC was performed. The average Log2FC of three replicates was plotted in a heat map. JQ1 attenuated PDGF and TGFB induced expression as early as 2 hrs for all targets, and was reduced to 4 targets by 16hrs. (**B)** The same targets were validated by immunoblot at 16 hrs and (**C)** quantified. Data are representative of 3 independent trials.

We also determined which targets were sensitive to JQ1 at the protein level **(Figure 3B)** and may therefore be relevant in altering contractile activity. The subset of targets included: Pdgfrα/β, αActn, Itgb1, Itga5, Src, and Cfl. Notably, PDGFRα/β, Src, Itga5 and Itgb1 decrease with JQ1 regardless of PDGF or TGFβ stimulation at 16 hrs. Additionally, we observed a reduction in the transcriptional expression of major contractile proteins such as Myh11, Sm22α and αSMA in both pHBSMCs and RBMCs **(Supplemental Figure 9A).** Additionally, Cnn1, but not Sm22α was found to be reduced at the protein level **(Supplemental Figure 9B)**. This further suggest that the contractility of both cell types may be sensitive to JQ1.

### JQ1 alters the cytoskeleton and inhibits contraction in SMC and fibroblasts

To determine whether the changes in expression of cytoskeleton-associated genes and proteins observed with JQ-1 treatment altered cell behavior, we determine how JQ1 impacts cytoskeletal and contractile changes induced by PDGF and TGFβ. Both human and rat cells were assessed using fluorescence-based imaging of phalloidin to assess the actin cytoskeleton. PDGF and TGFβ altered the cytoskeleton of both human and rat cell lines compared to baseline. PDGF induced a collapse-like phenotype whereas TGFβ increased actin fiber formation as reported previously by our group and others [7, 33-35], with JQ1 preventing these changes in both cell types. The most notable difference in both cell types was observed between TGFβ-treated cells in the absence and presence of JQ1, with RBMC exposed to JQ1 + TGFβ showing > 6-fold reduction in phalloidin intensity compared to RBMC treated with TGFβ1 alone (0.89e07 AU for TGFβ + JQ1 vs 6.02e07 (AU) for TGFβ)**(Figure 4B)**. In pHBSMC JQ1 inhibited a 20% increase in phalloidin intensity induced by TGFβ compared to DMSO control (1.76e07 AU for TGFβ vs 1.43e07 AU for TGFβ + JQ1) **(Figure 5B)**. Another difference between the two cell types in response to TGFβ included cell shape changes. Here, cell shape is defined as the circularity of the cell, defined further in the methods. TGFβ stimulation of RBMC resulted in a 3-fold increase in the circularity compared to control DMSO + Veh. JQ1 inhibited this TGFβ stimulated shape change. Interestingly, TGFβ stimulated pHBSMC showed a 50% decrease in circularity compared to control. This decrease in circularity was enhanced by JQ1 + TGFβ combination treatment compared to the DMSO control (0.2 to 0.1 respectively) **(Figure 4B and 5B)**.

**Figure 4:**
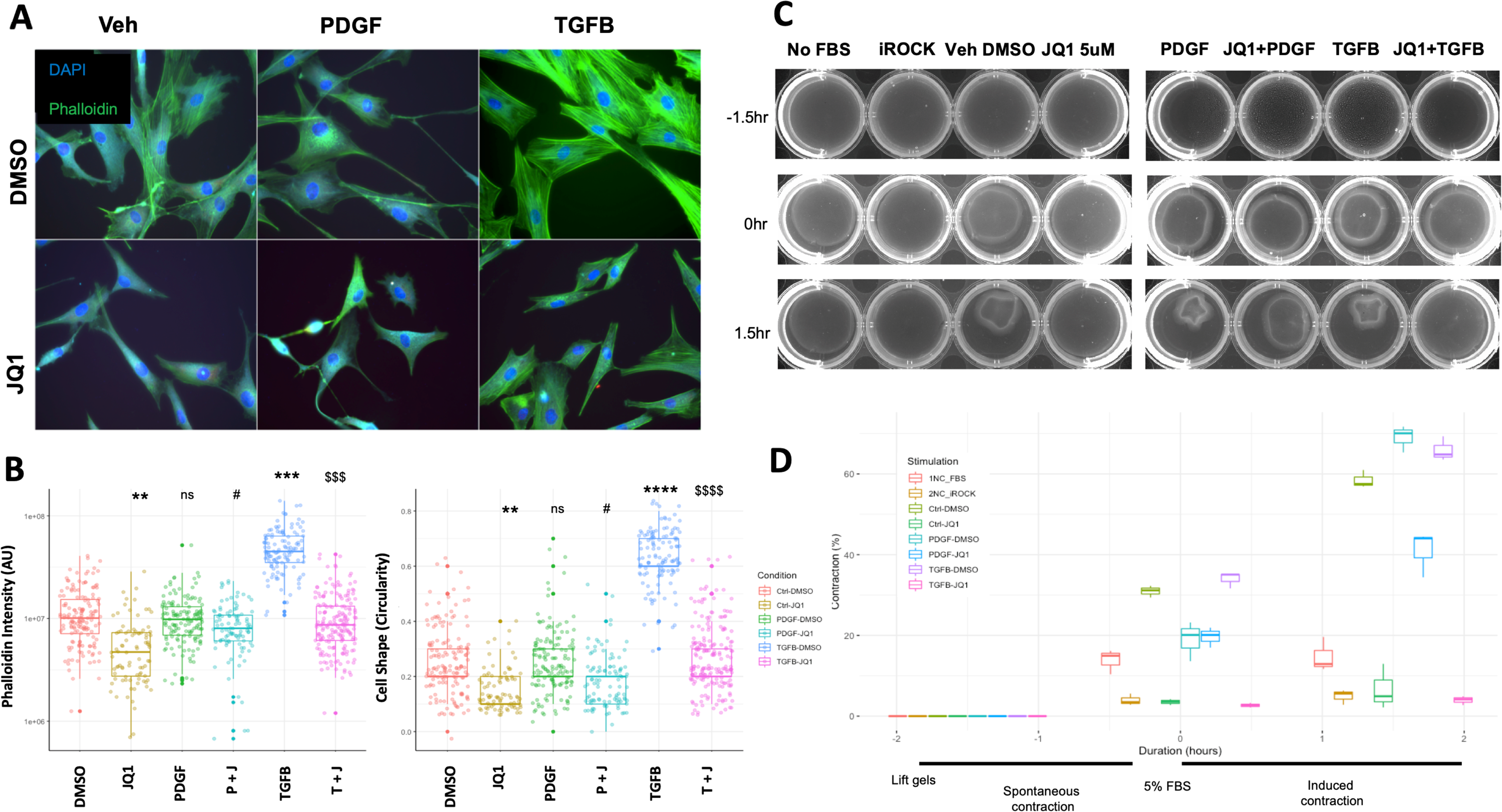
In vitro functional validation of JQ1 attenuation of PDGF and TGFB stimulated contraction in RBMC. **A)** Representative images of RBMC stimulated with PBS (Veh), PDGF, TGFB with DMSO or JQ1 for 16 hrs and stained with Vimentin, Phalloidin and DAPI. **B)** Phalloidin intensity and cell shape was quantified in ImageJ. **C)** RBMC were plated on 1.2 mg/mL collagen gels in a 12 well plate in 1mL of serum free media and incubated for 16 hrs with PBS (Veh), PDGF, TGFB with DMSO or JQ1. Select wells were stimulated for 30 mins with a ROCK inhibitor as a negative control for contraction. Collagen gels were separated from the walls of the wells and Imaged after 1.5 hrs to capture spontaneous (unstimulated contraction, followed by an additional 1.5 hrs in 5% FBS to facilitate contraction. **D)** Contraction was quantified using ImageJ and measured as a percent of the changed area of the gel from the baseline. Data are representative of 3 independent trials. *, p<0.05, ** p<0.01, ***, p<0.001, ****, p<0.0001 compared to control. #, p<0.05, ## p<0.01, ###, p<0.001, ###, p<0.0001 compared to PDGF. $, p<0.05, $$ p<0.01, $$$, p<0.001, $$$$, p<0.0001 compared to TGFB. Statistical significance was calculated with student ttest.

**Figure 5:**
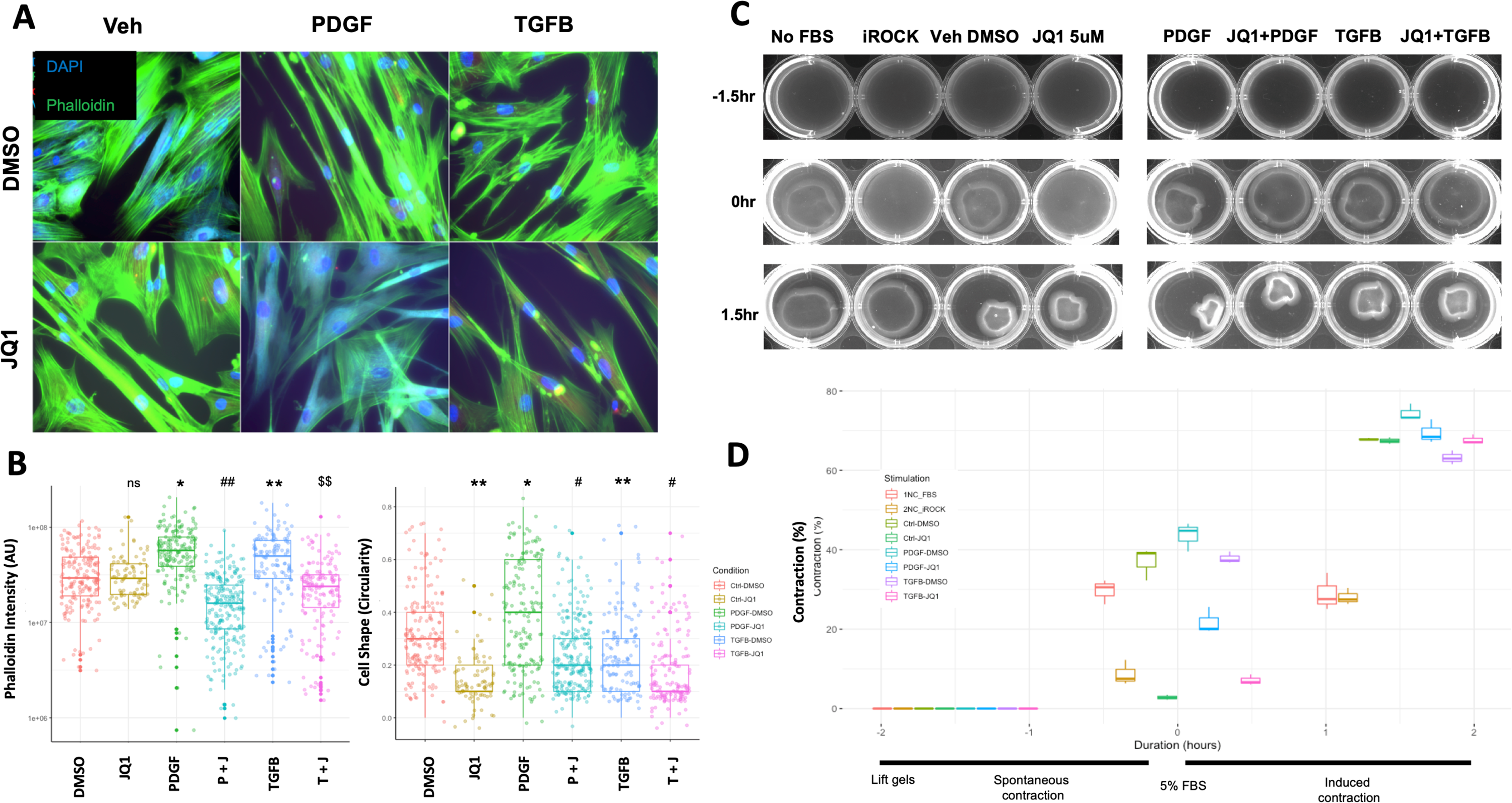
In vitro functional validation of JQ1 attenuation of PDGF and TGFB stimulated contraction in pHBSMC. **A)** Representative images of pHBSMC stimulated with PBS (Veh), PDGF, TGFB with DMSO or JQ1 for 16 hrs and stained with Vimentin, Phalloidin and DAPI. **B)** Phalloidin intensity and cell shape was quantified in ImageJ. **C)** RBMC were plated on 1.2mg/mL collagen gels in a 12well plate in 1mL of serum free media and incubated for 16 hrs with PBS (Veh), PDGF, TGFB with DMSO or JQ1. Select wells were stimulated for 30 mins with a ROCK inhibitor as a negative control for contraction. Collage gels were separated from the walls of the wells and Imaged after 1.5 hrs to capture spontaneous (unstimulated contraction, followed by an additional 1.5hrs in 5% FBS to facilitate contraction. **D)** Contraction was quantified using ImageJ and measured as a percent of the changed area of the gel from the baseline. Data are representative of 3 independent trials. *, p<0.05, ** p<0.01, ***, p<0.001, ****, p<0.0001 compared to control. #, p<0.05, ## p<0.01, ###, p<0.001, ###, p<0.0001 compared to PDGF. $, p<0.05, $$ p<0.01, $$$, p<0.001, $$$$, p<0.0001 compared to TGFB. Statistical significance was calculated with student ttest.

To determine if the changes we observed in the cytoskeleton that were attenuated by JQ1 had functional consequences we utilized an optimized and modified collagen gel contractility assay detailed in the methods.Contraction was assessed in the context of concurrent PDGF, TGFβ or vehicle (Veh) stimulation with JQ1 (5µM) or DMSO for 16 hrs. These parameters were determined in preliminary analysis as described in Methods. At baseline (Veh DMSO), both RBMC and HBSMC contracted collagen gels during both the spontaneous and elicited contraction phases **(Figure 4C and 5C)**. Spontaneous contraction was inhibited with the ROCK inhibitor Y-27632 (iROCK, 10 µM) in both cell types, albeit to a lesser extent in pHBSMC. Stimulation with PDGF and TGFβ for 16 hrs resulted in increased contraction above baseline for RBMC in the elicited contraction phase, which was attenuated with JQ1 pretreatment. In contrast, the magnitude of contraction by pHBSMC in the elicited contraction phase was not different whether cells were treated with PDGF, TGFβ or JQ-1. Cell viability was assessed in a dose-and time-dependent assessment of JQ1 on contraction using Alamar Blue **(Supplemental Figure 10 A, B**). JQ1 mediated inhibition of contraction was not due to cytotoxic effects.

### Inhibition of MYC dimerization destabilizes the cytoskeleton and reduces contractility similar to JQ1 in RBMC

JQ-1 inhibits MYC indirectly through targeting of the bromodomain-containing proteins BRD4 and BRD2 [36]. To further explore the impact of MYC inhibition on the cytoskeleton and contractile phenotype, we tested 2 additional MYC inhibitors with a distinct mechanism of action, namely the MYC-MAX dimerization inhibitors 10048-F4 (F4) and 10057-G5 (G5). Cytoskeletal changes were assessed using phalloidin staining. Cells were co-stained with DAPI and αSMA, as a marker of fibroblast activation **(Figure 6A)**. Phalloidin and αSMA intensity were quantified using ImageJ **(Figure 6B)**. All three MYC inhibitors reduced phalloidin intensity by more than 20%. αSMA signal, however, was increased by all inhibitors with the largest increases induced by F4 and G5 by 6-fold and 4-fold respectively. This different response likely reflects the direct and indirect mechanism of MYC inhibition by dimerization inhibitors versus JQ1.

**Figure 6:**
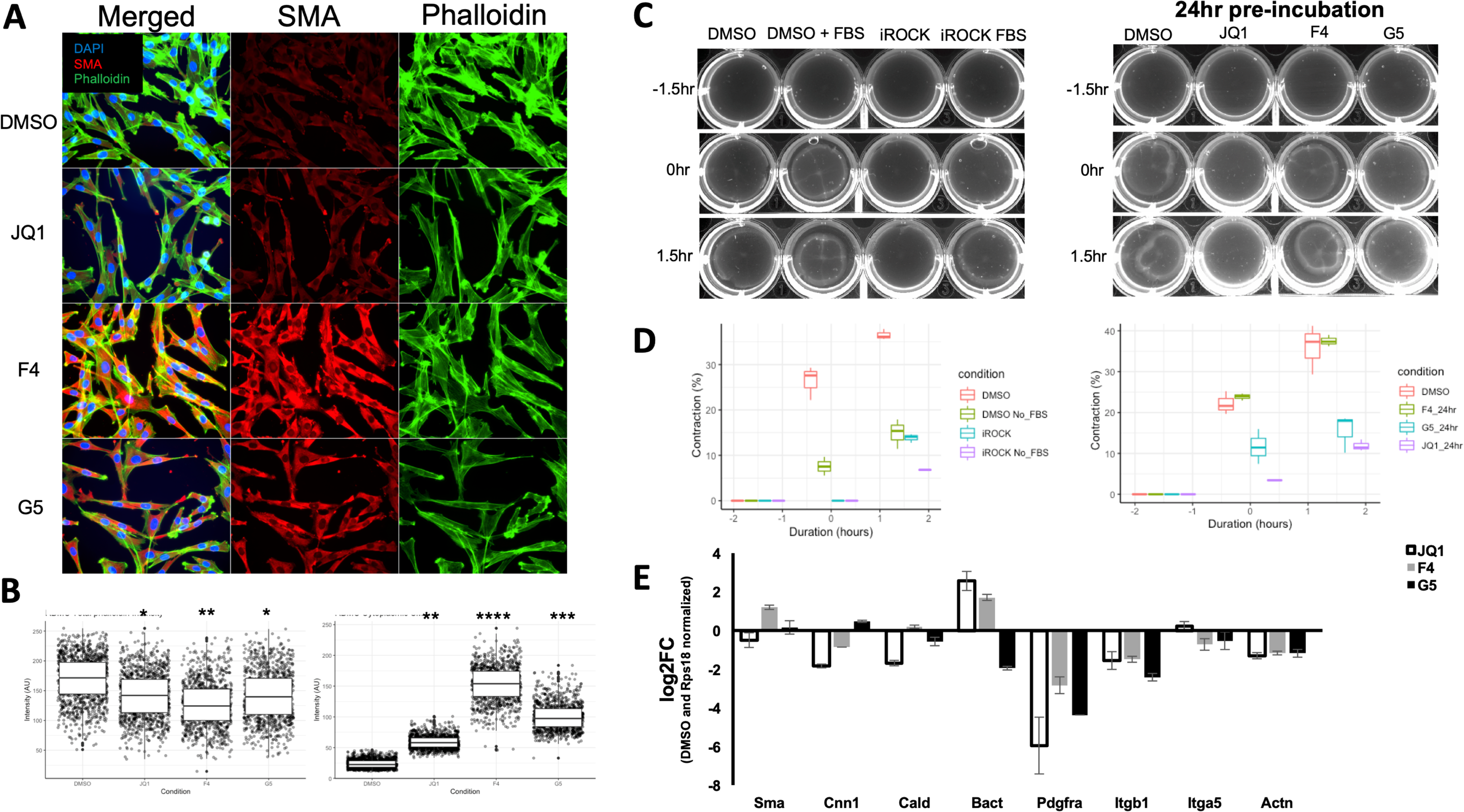
Inhibition of MYC–MAX dimerization destabilizes the cytoskeleton and reduces contractility similar to JQ1. RBMC were plated in 4 well chamber slides and incubated with DMSO, JQ1, MYC-MAX dimerization inhibitors 10048-F4 (F4), or 10074-G5 (G5) for 16hrs. Cells were stained for αSMA, Phalloidin or DAPI. Forty images were taken and stitched together with ImageJ to make a 5×8 field of view. **B)** Phalloidin and αSMA intensity were quantified for each cell using ImageJ and plotted for each condition. **C)** Cells were plated on collagen gels and stimulated for 16hrs with DMSO, JQ1, F4 or G5 in serum free media and spontaneous contraction of gels was captured for 1.5hrs prior to adding FBS to a final concentration of 5% to elicit contraction. **D)** Contraction was quantified as a percentage in the change in area of the gel compared to baseline. **E)** RNA was harvested from cells stimulated with DMSO, JQ1, F4 and G5 for 16hrs and predicted JQ1 sensitive cytoskeleton associated genes was measured with qPCR.

In the collagen gel contraction assay, JQ1 and G5, but not F4 were able to inhibit contraction from 21.1% in the DMSO control to only 4.6% and 11.4% during the spontaneous contraction phase, respectively. JQ1 and G5 were also able to inhibit elicited contraction from 37.2% with DMSO to 10.8% and 18.1% respectively. **(Figure 6C, D)**. The inhibition of RBMC contraction was coupled with reduced expression of cytoskeleton-associated genes as determined by qRT-PCR. All three inhibitors decreased expression of Pdgfrα, Itgb1 and Actn1 **(Figure 6E)**. Interestingly, there was no consensus on shared changes in smooth muscle contractile gene expression. However, JQ1 consistently down regulated αSma, Cnn1, and Cald1.

## Discussion

In this study, we build upon our prior demonstration of a MYC-centric network in SMC that regulates proliferation, to show that inhibition of MYC also attenuates contraction. We show that (i) PDGF-and TGFβ1-induced signals converge on MYC/MAX in smooth muscle cells and fibroblasts; (ii) JQ-1 treatment attenuates the expression of cytoskeleton-associated genes; and (iii) inhibition of MYC with two independent inhibitors reduces cytoskeletal stability and contraction of mechanically active cells. Together, these findings suggest that JQ-1 could be used to moderate aberrant contraction in addition to pathological fibroproliferative remodeling in hollow organs.

Although the ability of JQ-1 to target MYC activity has been most widely studied in the context of cancer [37], a number of recent studies have investigated the utility of JQ-1 in non-cancer settings including contractile tissues. Lim and colleagues evaluated the ability of JQ-1 to inhibit myometrial cell contraction in vitro and in vivo. In that study, either JQ1 treatment or knockdown of BRD2/3/4 were found to inhibit TNFα-stimulated adhesion and contraction of myometrial cells. JQ1 also inhibited inflammation-induced premature labor contractions in mice [38]. Notably, however, JQ1 was found to elicit its effects through inhibition of NFkB transcriptional regulation. Importantly the role of MYC was not assessed in that study. Similarly, Duan et al. showed that JQ-1 treatment could attenuate deleterious functional changes, including reduced left ventricle ejection fraction, in mouse models of cardiac injury [39], also through attenuation of NFkB signaling networks. In contrast, Yan and colleagues showed that JQ-1 could inhibit contractile responses in mouse aorta, rapidly and at high doses [40]. Moreover, a similar effect was observed with both (+)-JQ1 and its enantiomer (-)-JQ1 at 100µM, the latter of which cannot inhibit BRD4 recognition of acetylated histones [24]. These observations together with the demonstration that the inhibitory effect of (+)-JQ1 on smooth muscle contraction could not be replicated with knockout of BRD4, suggesting that the observed effect of JQ1 in that study reflected off-target activity. In contrast to Yan et al. short incubations with JQ1 in our analyses did not affect spontaneous or elicited contraction. “moreover, we observed that higher doses of 20uM resulted in cytotoxicity”. (**Supplemental Figure 9).** Our studies do support a mechanism for aberrant contractile inhibition by JQ1 through the repression of the MYC and likely Jun TF regulator gene networks.

Previous studies have established a connection between MYC and contraction through interactions with an oncogenic form of RHOA. However, this relationship has been shown to down regulate the F-actin stress fiber formation [13, 15]. Additionally, overexpression of MYC alone has been shown to be sufficient to destabilize the actin cytoskeleton [17]. Notably, these studies attempted to recapitulate a cancer like state though over expressing MYC, which does not reflect the physiologically relevant stimulation of MYC. Our data demonstrate that MYC inhibition destabilizes the actin cytoskeleton. To address this dichotomy, we compared the transcriptomic changes of fibroblasts forced to overexpress MYC with that of proliferative fibroblast (Supplemental Figures 13-15) using publicly available data. While there was significant overlap in DEGs induced by both stimulations, analysis of MYC induced DEGs were associated with RNA and DNA transcriptional regulation while down regulated genes were associated with actin cytoskeleton regulation. Importantly, proliferation induced DEGs were associated with actin cytoskeleton regulation while down regulated genes were enriched for RNA binding and histone acetyltransferase activity. The JQ1 mediated inhibition of physiologically relevant stimulation of MYC that we have demonstrated combined with MYC overexpression studies supports a complex role for MYC expression dynamically influencing cytoskeletal rearrangement and stabilization in mechanically active cells in addition to its well-studied oncogenic effects in other cell and tissue types [41-44].

Prior findings from us and others have demonstrated opposing effects of PDGF and TGFβ on SMC, with the former acting as a canonical mitogen and the latter, a potent growth inhibitor [22, 45]. Interestingly, however, in silico analysis of publicly available data revealed significant overlap in the genes differentially regulated by PDGF and TGFβ in both SMC and fibroblasts, with more than 80 DEGs regulated by MYC, MAX, and JUN in master regulator analysis. Comparative analysis between our previously published data of PDGF stimulated pHBSMC showed nearly 30% similarity with datasets of fibroblasts and smooth muscle cells stimulated with PDGF or TGFβ1. These observations are in agreement with data from Ghosh et al., which established that PDGF and TGFβ shared significant overlap of differentially expressed genes within mesenchymal stem cells [46]. In that study, the authors proceeded to compare cells stimulated with each cytokine alone to those exposed to both concurrently, adding that the combination influenced cell stiffness through synergistic induction of integrin expression. Notably, the genes encoding α5 and β1 integrins (Itga5 and Itgb1) were induced in the datasets we analyzed and attenuated with JQ1 *in silico* and *in vitro*. Consistent with their role in anchoring cells to their substrate, the reduction in or loss of mediators of cell-ECM connections, such as integrins, has been shown to alter contraction [47, 48]. Our data support a model in which JQ1 reduces aberrant contraction, at least in part through repressing MYC-mediated regulation of Itga5 and Itgb1.

It has been shown that aberrant contraction is a consequence of alterations to the cytoskeleton of mechanically active cells [2, 49]. We show that JQ1-induced destabilization of the cytoskeleton results in reduced contraction. Interestingly, These findings agree with those of Stratton et al. who showed that JQ1 reduced baseline and TGFβ1-stimulated cytoskeletal stabilization of cardiac fibroblasts, with a corresponding decrease in collagen gel contraction [31]. In that study the authors did not address the role of MYC. Instead NFkB was found to mediate this process. In our study, we observed that the effect of JQ1 could be replicated with another MYC inhibitor with a different mechanism of action. Similarly, Wang and colleagues identified that JQ1 and the MYC inhibitor 10048-F4 repressed MYC expression in rat and rabbit lens epithelial cells [50]. This resulted in reduced EMT as manifested by αSMA and fibronectin expression further supporting a MYC dependent phenomenon of JQ1 inhibition. Our data support that the inhibitory effect of JQ1 on aberrant contraction is being mediated through inhibiting MYC in mechanically active cells.

While this study provides evidence of a MYC centric network of aberrant gene expression in smooth muscle cells and fibroblasts stimulated by PDGF and TGFβ, these comparisons have been limited to *in silico* and *in vitro* analyses. As mentioned previously, there have been several studies that have utilized JQ1 to attenuated pathological changes in various animal models of organ injury [20]. However, most have not implicated or compared MYC activity and inhibition in the mechanism of action for JQ1 in the attenuation of those changes. Our data support a potential for repurposing the anti-cancer, therapeutic JQ1 to inhibit MYC in a non-cancer setting. However, understanding the extent to which MYC expression and activity can be different in these contexts will provide further rationale for its efficacy. Finally, when considering the primary target of JQ1 activity, BRD4, which prevents MYC recruitment and transcriptional activity, there are other BRD proteins that JQ1 can inhibit. Therefore it is important to understand the extent to which the effects we have observed by JQ1 are BRD4 and MYC specific or are off-target effects.

## Conclusion

Utilizing in silico and in vitro techniques we determined that the MYC inhibitor, JQ1, attenuates MYC-driven, aberrant cytoskeletal changes and contraction induced by PDGF and TGFβ1 treatment of smooth muscle cells and fibroblasts. Our findings support a shared, MYC-regulated, gene network stimulated by PDGF and TGFβ1 in mechanically active cells. We showed that JQ1 attenuates aberrant contraction and cytoskeletal changes through regulating the expression of various cytoskeleton associated genes. Further, comparing the effects of MYC-Max dimerization inhibitors with JQ1 suggest that there are partially overlapping gene targets between BRD4 and MYC-MAX transcription factors that result in similar impacts to the cytoskeleton and contraction.

## Methods

### In silico analyses of publicly available microarray data

We re-analyzed gene expression data generated previously by us [8] and compared the results with re-analyzed publicly available data of SMC and fibroblasts stimulated with PDGF and TGFβ1. Data were retrieved from the NCBI GEODatabase using GEOQuery [51]. Three datasets were retrieved; GSE63383 (SMC stimulated with TGFβ), GSE61128 (Fibroblasts stimulated with TGFβ), and GSE14256 (SMC stimulated with PDGF). Data quality was assessed using PCA plots to determine how replicates of various stimulations clustered together using the base R stats package [52]. Well-clustered replicates that were separated from the other stimulations within a dataset were analyzed using LIMMA based packages in R [53]. To perform enrichment analysis, DEGs from each comparison as well as Log2 fold changes and p-values were incorporated into the pathfinder package in R [54]. Separate analyses were run using GO terms for molecular function, biological process and for cellular compartment.

### In silico analyses of publicly available RNA-seq data

We reanalyzed publicly available data which included rat VSMCs exposed to PDGF (GSE111714), human VSMC stimulated with undefined growth medium (GSE138323), rat cardiac fibroblasts stimulated TGFβ (GSE127229)[31] and Rheumatoid Synovial Fibroblasts stimulated with IL1B (GSE148395)[32]. All datasets contained conditions of JQ1 with and without the respective stimulus. For GSE111714, GSE148395, and GSE127229, SRA raw files were downloaded from the SRA run selector using the SRAtoolKit [55], genome indexes were generated using the most recent versions of the appropriate species primary assemblies, aligned to the appropriate species library using STAR aligner [56], count matrices were generated using the Subreads, FeatureCount function [57] and the count matrices were analyzed with DESeq2 [58]. Data frame manipulations were performed using a variety of packages in R including tidyR [59], dplyR [60], tibble [61], and stringR [62]. Plots were constructed using ggplot2 [63] and GGally [64]. Conversion of Ensembl and Affymetrix microarray gene IDs was performed using a combination of biomaRt [65], and Hs.eg.db [66].

Venn diagrams of overlapping genes were generated with ggVennDiagram [67] and Volcano plots were generated using EnhancedVolcano [68]. Pattern analysis of DEGs was performed using the DEGpattern [69] package. ClusterProfileR [70] and pathfindR [54] were utilize to perform enrichment analysis using GO terms for molecular function, biological process and cellular compartment on DEG lists without and with Log2FC and adjusted pvalues, respectively. Hierarchical clustering of enriched terms was performed to cluster similar groups of terms together.

### ChEA3 Transcription factor enrichment analysis

The ChEA3 transcription factor (TF) enrichment analysis tool [23] was used to identify and rank TFs that regulate the list of DEGs from each comparison. The top ten TF identified using a list of DEGs was generated using the ENCODE database. Bar charts showing the -log Fisher’s exact test (FET) p-value were generated for the top ten TFs.

### Cultured cells

Primary human bladder smooth muscle cells were purchased from ScienCell and cultured in Smooth muscle cell medium (ScienCell) with Penicillin/streptomycin, 2% FBS and Smooth muscle cell growth factors (ScienCell). Cells were used up to passage 6. Rat bladder mesenchymal cells were isolated from P10 rat pups as follows: Twenty bladders were minced in 1 ml of 1x PBS post dissection and incubated for 1 hour in a dissociation solution containing: 1.25 mg/ml Elastase, type III 1 (E0127, 20mg, Sigma), 1 mg/ml Collagenase I (C0130, 1 g, Sigma), 0.25 mg/mL Trypsin Inhibitor (soybean, T9003, 100 mg, Sigma), BSA, 2mg, and 0.2 mL of pen/strep (100 X) in 20 mL of M199 media (11150059, 500 mL Thermo Fisher). Following incubation, the dissociated cell suspension was filtered through a 100 µm filter, centrifuged at 176 x g for 3 mins, resuspended in 10 mL of M199 containing 20% FBS and plated in a 10 cm dish. Cultures were passaged 4 times before cells spontaneously immortalized. RBMC were used at passages 16-20.

### Validation of *in silico* predicted JQ1 sensitive genes using qRT-PCR

Seventeen cytoskeletal targets were predicted to be sensitive to JQ1 after comparative analysis of the three sequencing datasets generated from mesenchymal cells stimulated with a cytokine and treated with JQ1. Seven of the seventeen targets were validated using qPCR and Immunoblots. Primers to all seven targets were designed for human and rat transcripts using NCBI primer design tool such that the length of the product was between 100-300 bases to facilitate rapid detection via qPCR and ordered from Integrated DNA technologies. RBMC and pHBSMC were plated in 6 cm dishes at 70% confluence in complete media (DMEM, 10% FBS, pen/strep). After an overnight incubation, the media was changed to low serum media (DMEM, 0.5% FBS, pen/strep) (Thermo Fisher) for 24 hrs. Cells were then stimulated with DMSO (Sigma Millipore), PBS (Thermo Fisher), JQ1 (5 µM) (Cayman Biochemical), PDGF (2.5 ng/mL) (R&D systems) or, TGFβ1 (2.5 ng/mL) (R&D systems). Cells were harvested after 24 hrs using 500µL of TRIzol. 100µL of chloroform was added to the Trizol, mixed by vortexing and centrifuged for 15 mins at 7826 x g. The aqueous phase was separated and added to an equal volume of 70% ethanol before using the RNeasy minikit (Qiagen) to isolate RNA following the manufacturer’s protocol. cDNA was generated using the iScript cDNA synthesis kit (Bio-Rad) following the manufacturer’s protocol. qRT-PCR was performed using 18ng of sample cDNA with 1uL of premixed primers and 10uL of 2x SYBR select master mix (Thermo Fisher) and run in a QuantStudio3 thermocycler (Thermo Fisher). Expression of each target was normalized to Gapdh, Rps18 as housekeeping genes with the least variation in cycle values across all samples. Notably, BACT could not be used as a house keeping gene as the standard deviation across samples was greater than 1 cycle and had a range of 4.78 cycles, while GAPDH and Rps18 had standard deviations of 0.742, and 0.148 respectively and a range in cycle values of 3.04 and 0.59 respectively (data not shown)

### Validation of *in silico* predicted JQ1 sensitive genes via immunoblots

Targets assessed via immunoblot included PDGFRβ, PDGFRα, ITGB1, ITGA5, αACTN, SRC, and CFL (Cell Signaling Technology). Additional targets assessed included MYC, SM22α (Cell Signaling Technology), CNN1, Vim, αSMA and BACT (Sigma). To assess these targets via immunoblot, cells were stimulated as before in 10cm dishes, washed with 1x PBS and then lysed on ice using 1x cell lysis buffer (Cell Signaling Technology) containing SDS. The lysate was mixed with 4x sample buffer (1% Bromophenol blue, 50% glycerol, 0.125 M Tris-HCl, pH 6.8, 4% SDS) and boiled for 15 mins. Lysates were resolved on 10% and 15% polyacrylamide mini gels at 100V for 20 mins then 160V for 70 mins. Gels were transferred to nitrocellulose membranes at 100V for 2 hrs at 4°C. Nitrocellulose membranes were stained with Ponceau S to confirm transfer of total protein then washed with distilled water for 5mins, PBS-T for 5mins and blocked in 10% milk for 1 hr. Membranes were then washed briefly with PBS-T and incubated in one of the aforementioned primary antibodies overnight at 4°C. Membranes were then washed 3 times for 15 mins in PBS-T prior to being incubated in 2° HRP conjugated anti-rabbit or anti-mouse antibody for 1 hr at room temperature in 10% non-fat powdered milk followed by 3 washes in PBS-T. The washed membranes were then subjected to 5min incubation with SuperSignal™ West Pico PLUS Chemiluminescent Substrate (Thermo Fisher) following the manufacturers protocol. Signal was captured by ChemiDoc chemiluminescence imaging or on HyBlot CL™ Autoradiography Film (Thomas Scientific) after 10 sec, 1 min, 1 hr, 1 min, 30 sec, 10 sec exposures. Films were digitized using an Epson scanner and quantified with FIJI.

### Assessment of cytoskeleton changes

pHBSMC and RBMC were plated into 2x 4 well chamber slides (Thermo Fisher) at a concentration of 50k cells/well in complete medium. Following an 8 hr incubation at 37°C and 5% CO_2_ the media was changed to DMEM with 0.5% FBS and incubated for 24 hrs. Cells were stimulated with DMSO, PBS, JQ1, PDGF, TGFβ1, JQ1 + PDGF, JQ1 + TGFβ1 for 24 hrs followed by fixation using 4% paraformaldehyde for 15 mins at room temperature. Post fixation, cells were blocked and permeabilized using blocking buffer (5% FBS, 0.1% BSA, 3% Triton X in PBS) for 1hr at room temperature. Permeabilized cells were then incubated overnight in Alexa Fluor 488 phalloidin (Thermo Fisher), anti-αSMA, anti-MYC. The cells were washed in 1 x PBS 3 times for 5 mins and then incubated in incubation buffer (1% FBS, 0.1% BSA in PBS) for 1 hr containing species specific 2° Antibodies conjugated to Alexa Fluor 594 or Alexa Fluor 647 (ThermoFisher). The cells were then washed again, and a cover slip was added after addition of DAPI containing mounting medium (Vector Labs). The cells were imaged using a Zeiss fluorescence scope. 20-30 images were taken for each condition to ensure that more than 100 cells were captured per condition. Fluorescent signal and cell shape was quantified via FIJI based macro. The circularity of the cell is calculated as the ratio of the second longest dimension of the cell that runs perpendicular to the longest dimension of the cell. The closer the ratio is to 1 the more rectangular the cells. Long thin cells will have a circularity closer to 0.

### Assessment of changes in contractility

Contractility was assessed using a modified collagen gel contractility protocol. Briefly, Collagen gels were made using 2.4mL 10x PBS, 157uL of 1N NaOH, 14.07mL of ddH_2_O, 6.56mL of rat tail type 1 collagen (Advanced Biomatrix), all reagents were kept on ice until ready to mix. Once mixed, 0.75µL of the solution was added to every well of 2 x 12 well plates. Gels were incubated at 37°C in 5% CO_2_ for 1hr to polymerize. Cells were then lifted, counted, spun down to remove trypsin containing media, and resuspended in DMEM with 0.5% FBS and Pen/Strep at a concentration of 200k cells/mL. 1mL of cell suspension was added to each well containing a solidified collagen gel. Cells were incubated overnight at 37°C in 5% CO_2_. Cells on gels were then stimulated for 16 hrs with DMSO, ROCK inhibitor Y-27632 (iROCK), PDGF, TGFβ1, JQ1, PDGF + JQ1, or TGFβ1 + JQ1. Gels were then separated from walls and bottom of the well and imaged every 30 mins for 1.5 hrs using a chemi-doc imaging platform (Bio-Rad). This first 1.5 hrs was considered the spontaneous contraction phase (SCP). After the SCP, 100uL of FBS was added to each well containing cells on gels to initiate the elicited contraction phase (ECP). 16 hr pretreatment with 0.5µM JQ1 led to modest inhibition of contraction, while 5.0µM JQ1 lead to significantly inhibited contraction with limited reduction in viability. All doses of JQ1 at 24 hrs reduced viability by nearly 30%. As a result, 5µM JQ1 pretreatment for 16 hrs was used for all additional collagen gel contraction assays unless otherwise noted Images were taken every 30 mins for 1.5 hrs and captured with the chemi-doc imaging platform. Images were analyzed in FIJI and the data was plotted in R using ggplot2.

## Supplemental Figure Legends

**Supplemental Figure 1: PCA plots of PDGF and TGFB stimulated SMC and Fibroblasts datasets**

4 microarray datasets were identified from NCBI GEOdatabase in which fibroblasts or smooth muscle cells were stimulated with PDGF or TGFB. PCA plots were generated using the raw data to determine the extent of variation between conditions and replicates. Of the 4 datasets, only GSE61128 did not pass the filtering criteria phase, which required greater than 20% of the variation to be attributed between the conditions.

**Supplemental Figure 2: Enrichment analysis using GO-CC terms; changes in chromatin remodeling and cytoskeleton**

Shared DEGs were identified between our PDGF stimulated pHBSMC dataset and each of the three other datasets. Gene ontology enrichment analysis and clustering of CC terms was performed. In all comparisons of our dataset, either with **A)** GSE63383 in which smooth muscle cells were stimulated with TGFβ, or **B)** GSE14256 in which fibroblasts were stimulated with PDGF, or **C)** GSE61128 in which fibroblasts were stimulated TGFβ, were enriched for changes in chromatin (highlighted with orange boxes) and for changes in the cytoskeleton (highlighted with green boxes).

**Supplemental Figure 3: Identification of shared genes between all 4 datasets, TF regulator analysis and enrichment analysis**

DEGs compared **A)** via a 4 way Venn-diagram to identify the number of genes that were common to all four datasets. **B)** MYC and MAX were among the top TF predicted to regulate these shared genes based on -log Fisher’s exact test (FET) p-value. Enrichment analysis of the shared genes was performed using **C)** GO-MF terms, **D)** GO-BP terms and **E)** GO-CC terms.

**Supplemental Figure 4: Quality control check of Data sets using PCA plots to determine homogeneity among treatment groups**

Quality control assessment of publicly available data in which mesenchymal cells are stimulated with PDGF (GSE111714), Growth media (GSE138323), IL1β (GSE14395), or TGFβ (GSE127229) with or without JQ1 using PCA plots. More than 80% of the variation for each dataset was between the conditions and not the replicates.

**Supplemental Figure 5: Enrichment analysis of JQ1 sensitive genes using GO-MF terms** Enrichment analysis and clustering of JQ1 sensitive genes using GO-MF terms. JQ1 sensitive genes from each dataset **A-D)** were enriched for Chromatin related changes (highlighted in orange) and cytoskeleton related changes (highlighted in green).

**Supplemental Figure 6: Comparison of cytoskeletal effectors to validate *in vitro***

**A)** A heatmap of the expression of cytoskeleton associated genes under cytokine and cytokine + JQ1 stimulation. **B)** A subset of the JQ1 sensitive cytoskeleton associated genes from the heatmap were overlaid onto the Regulation of Actin Cytoskeleton KEGG pathway. Genes induced by JQ1 are highlighted in red and down regulated genes are highlighted in blue. **C)** A table of JQ1 sensitive hits from the KEGG pathway.

**Supplemental Figure 7: HBSMC and RBMC comparison and JQ1 dose and time effect on cellular viability**

**A)** A heatmap of C_t_ values from qPCR data comparing RBMC and HBMC gene expression included ECM, smooth muscle and fibroblast specific gene expression. Higher expression is color coded in blue and lower expression is color coded in red. **B)** Dose and time response of RBMC stimulated with JQ1.

**Supplemental Figure 8: Validation of JQ1 sensitive cytoskeletal genes in pHBSMC**

**A)** A heatmap of Log2FC values from qPCR data assessing a time course of Veh, PDGF or TGFβ stimulation with or without JQ1. Data are representative of 3 independent trials.

**Supplemental figure 9: JQ1 robustly reduces transcription of contractile genes human and rat bladder contractile cells**

**A)** Smooth muscle markers were assessed in both the RBMC and pHBSMC via qPCR and **B)** immunoblots. MYC was also assessed via immunoblot to confirm JQ1 reduced MYC protein expression.

**Supplemental figure 10: JQ1 inhibits contraction of RBMCs on collagen gels after 16hrs with low cytotoxicity**

**A)** A dose and time response of JQ1 mediated inhibition of RBMC contraction of collagen gels and **B)** assessment of cell viability using Alamar blue. Bar charts represent the means three independent replicates. Error bars reflect the standard deviation. pvalues were calculated with student ttest. * pvalue < 0.05, ** pvalue < 0.01, *** pvalue < 0.001. Data are representative of 3 independent trials.

