## Supplemental Figures for "Investigation of the impact of bromodomain inhibition on cytoskeleton stability and contraction"

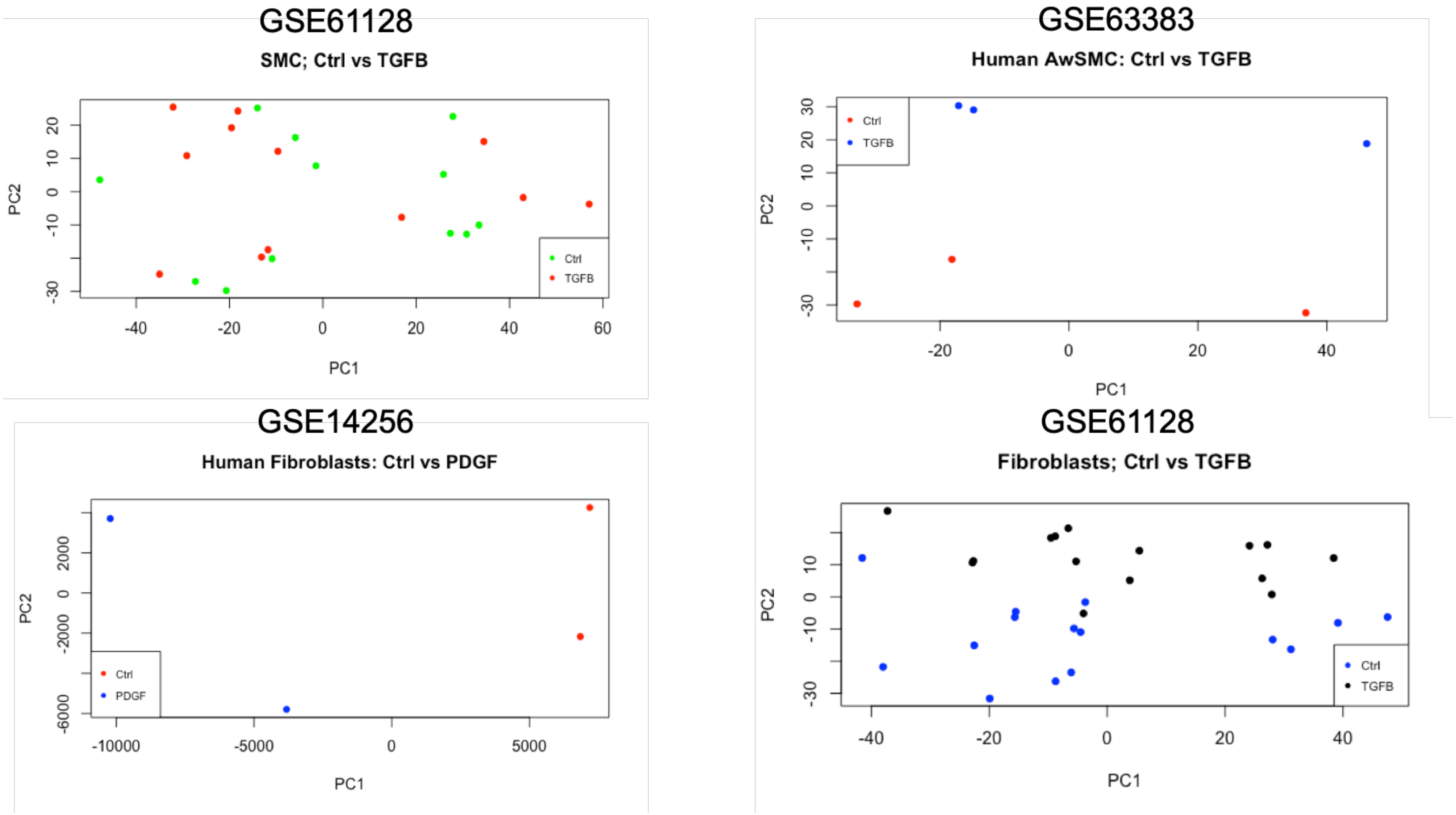


**Supplemental Figure 1: PCA plots of PDGF and TGFB stimulated SMC and Fibroblasts datasets**

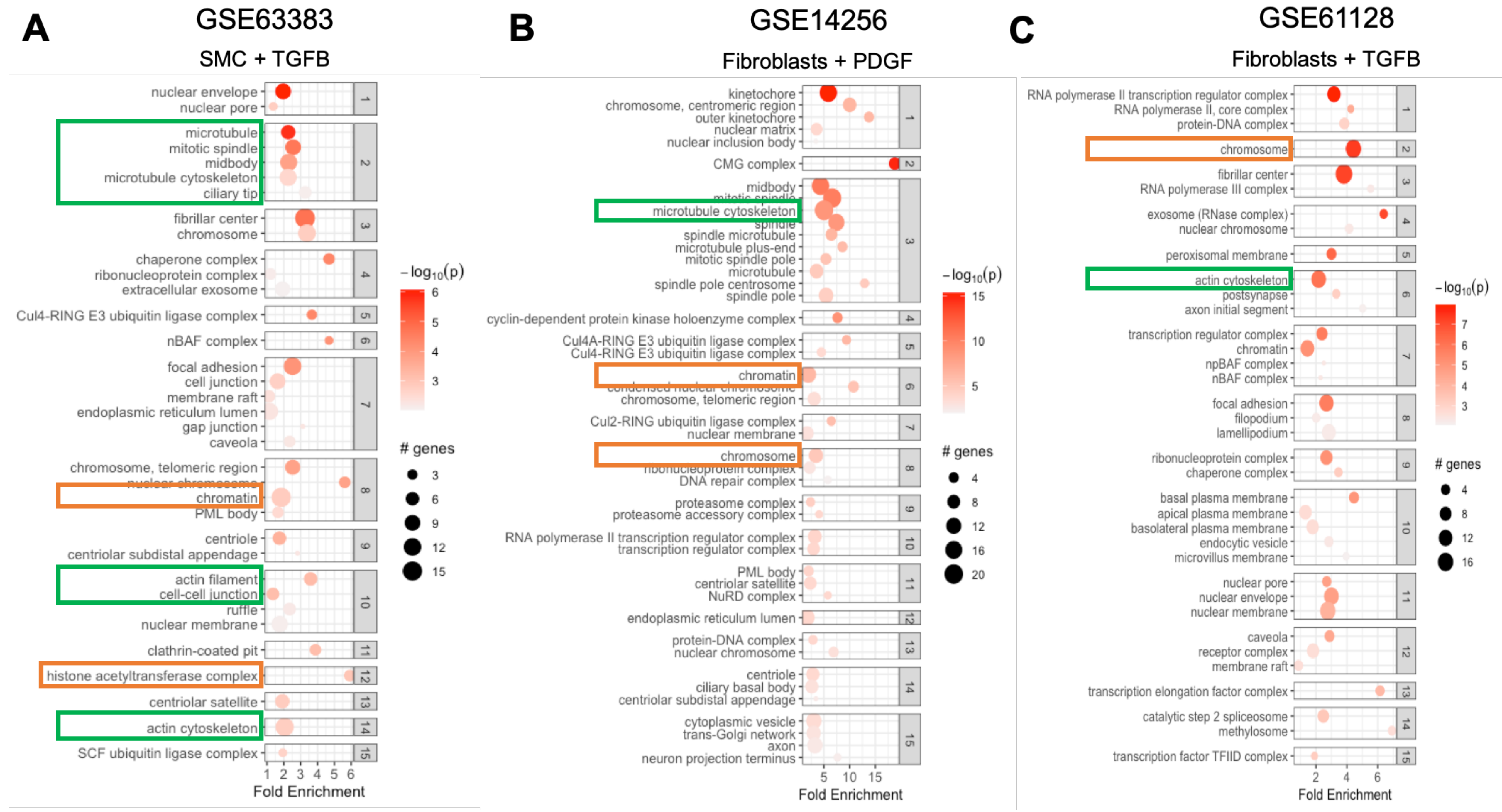


**Supplemental Figure2: Enrichment analysis using GO-CC terms; changes in chromatin remodeling and cytoskeleton**

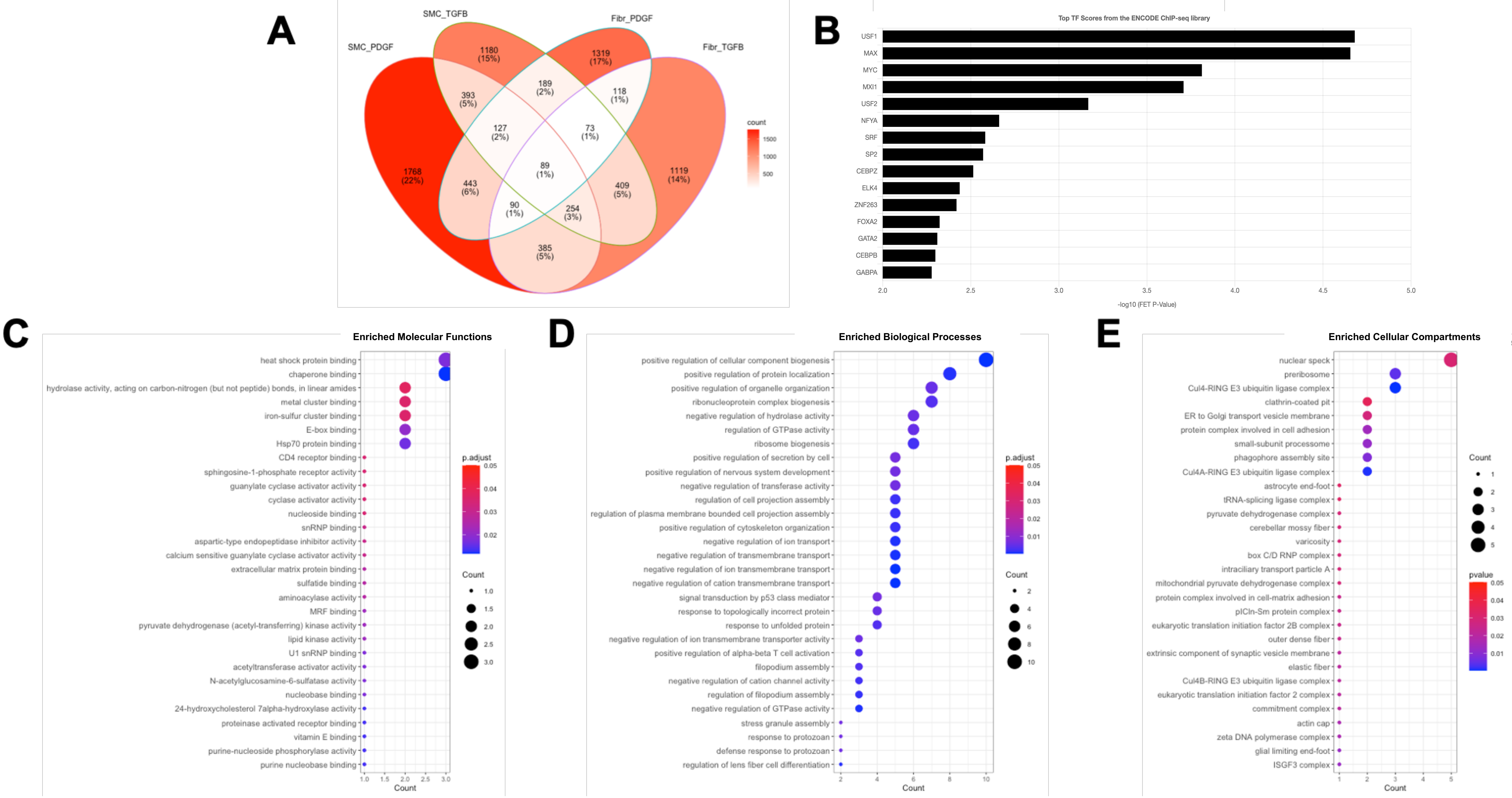


**Supplemental Figure 3: Identification of shared genes between all 4 datasets, TF regulator analysis and enrichment analysis**

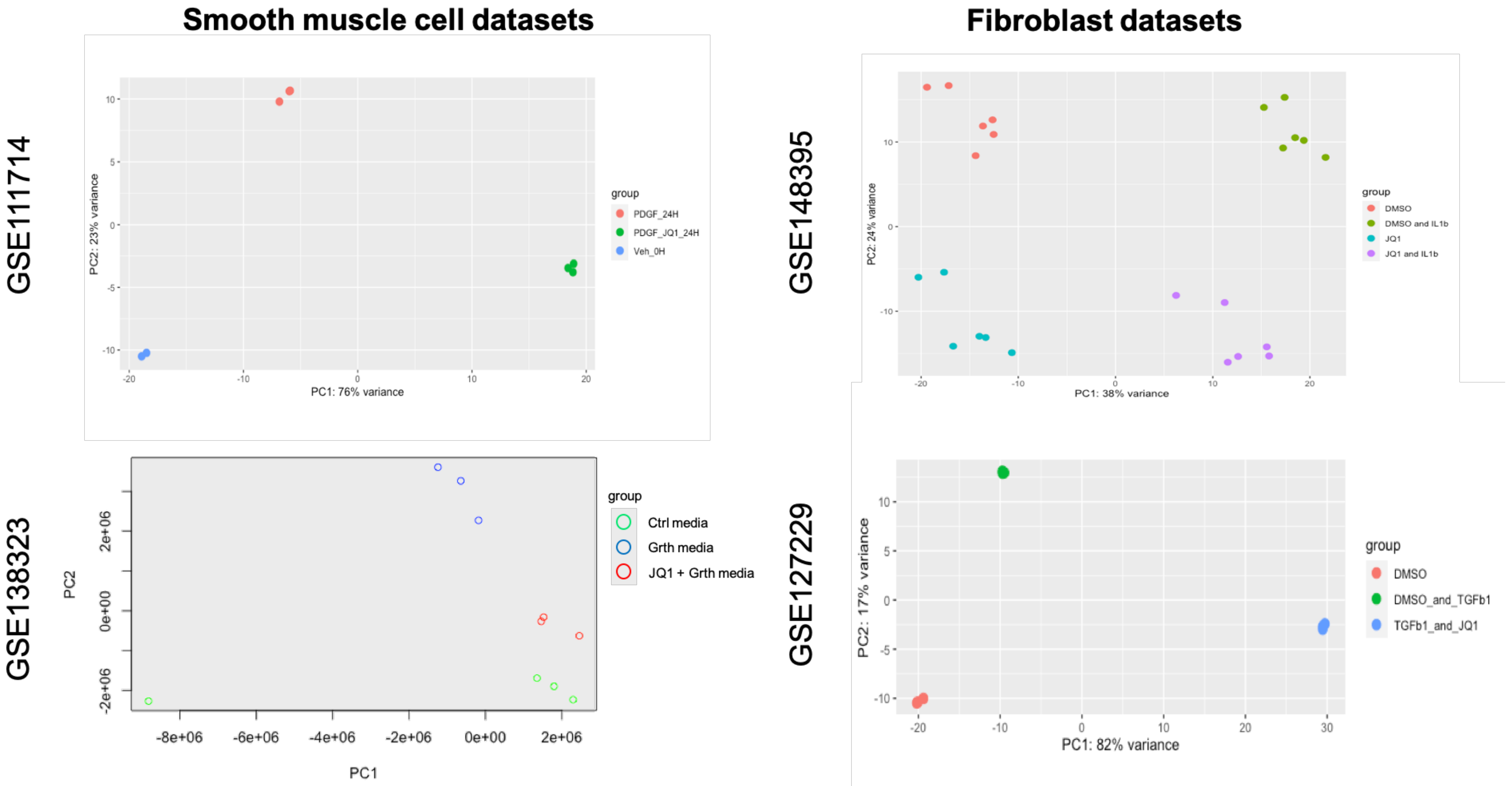


**Supplemental Figure 4: Quality control check of Data sets using PCA plots to determine homogeneity among treatment groups**

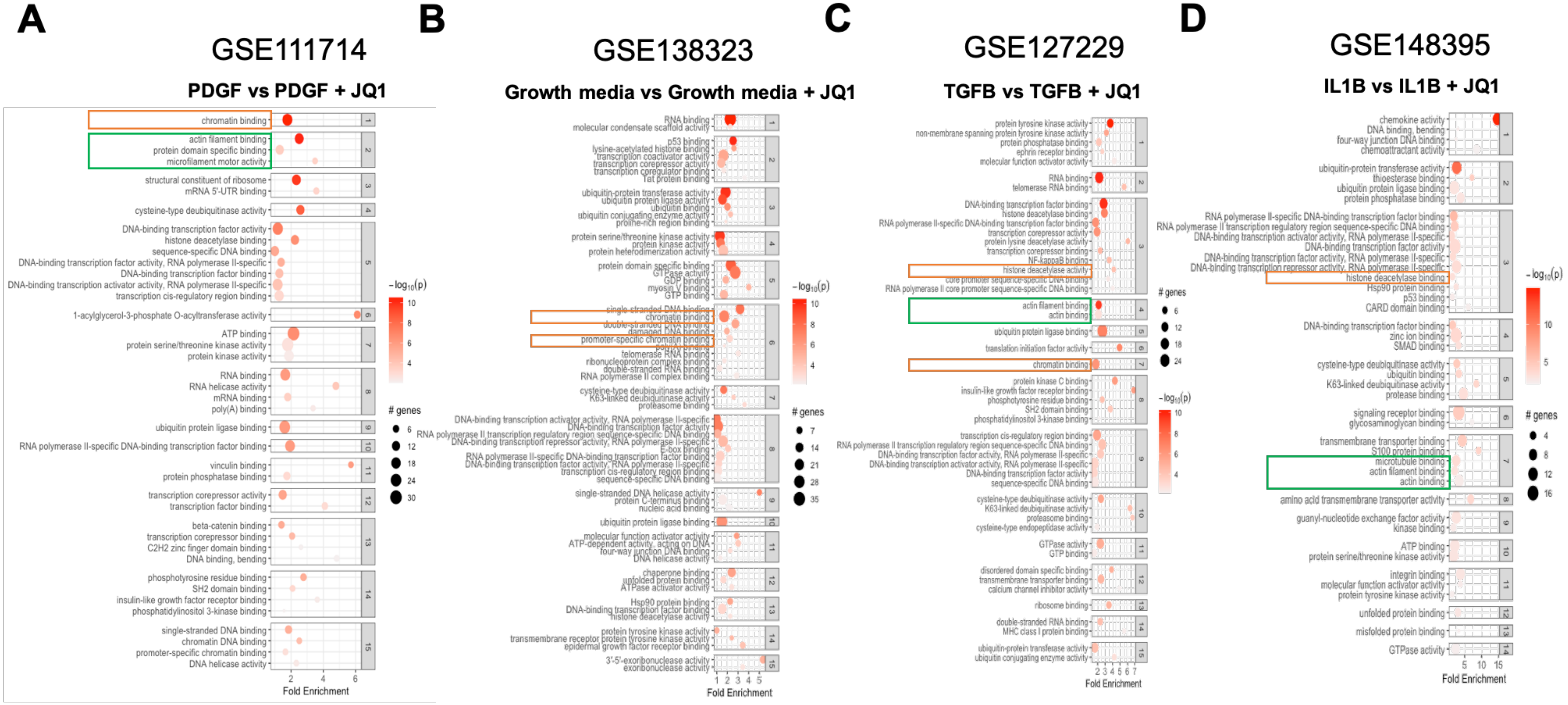


**Supplemental Figure 5: Enrichment analysis of JQ1 sensitive genes using GO-MF terms**

A

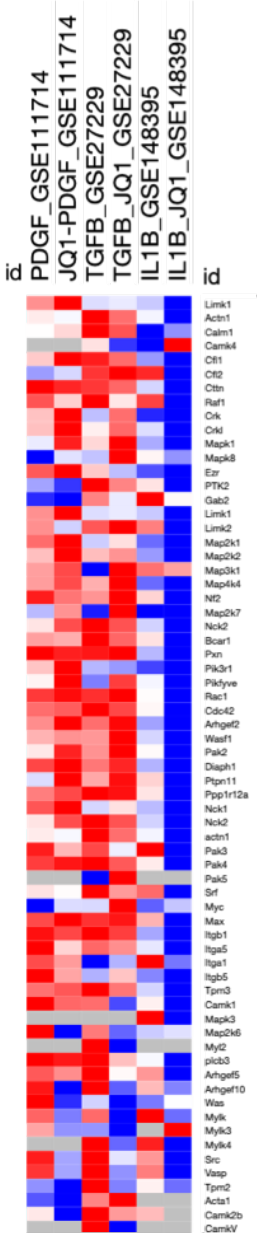


B

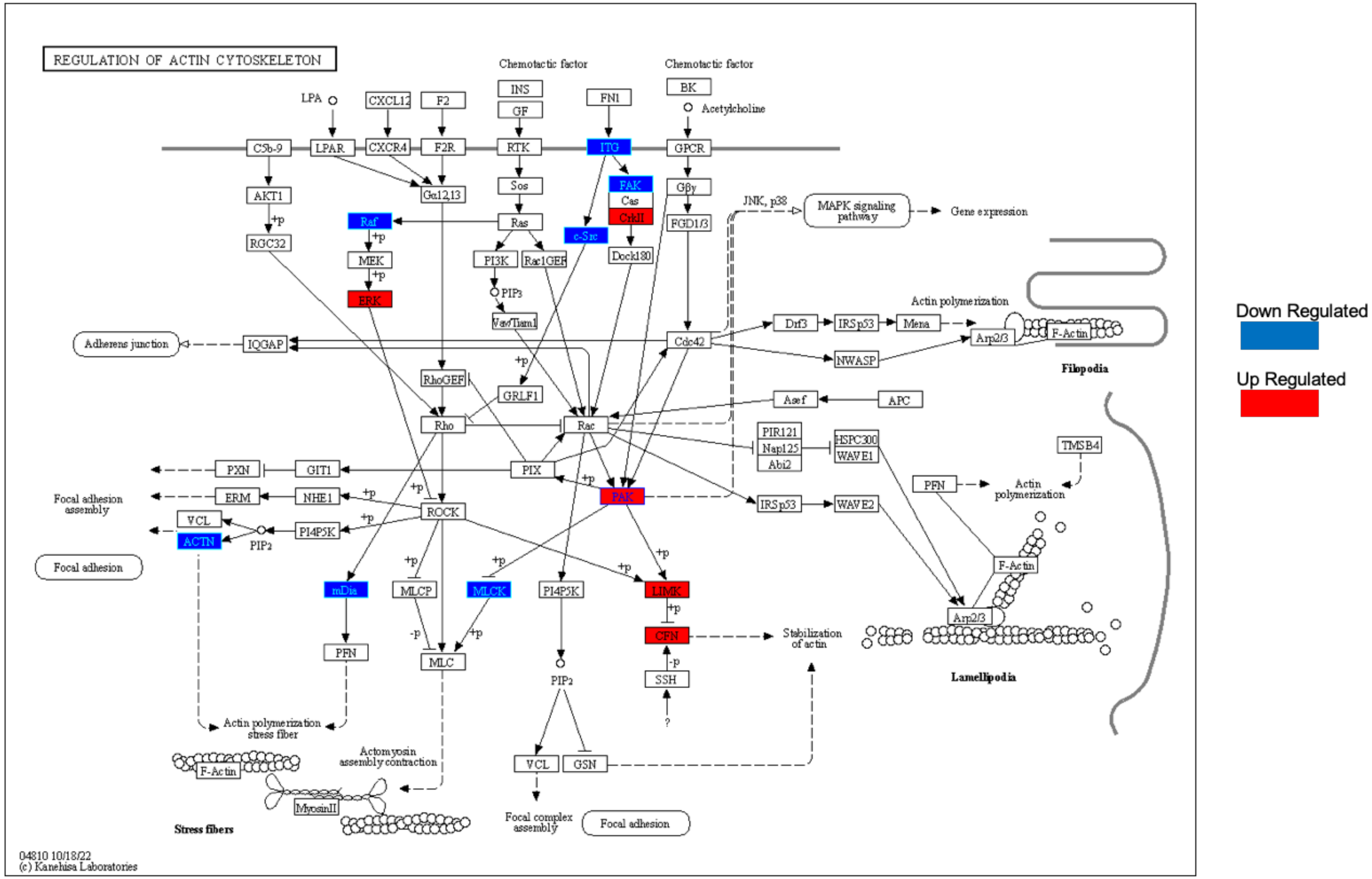


C

| Predicted JQ1 target genes | |
|---|---|
| Raf1 | Raf-1 Proto-Oncogene, Serine/Threonine Kinase |
| PTK2 | Protein Tyrosine Kinase 2 / Focal adhesion kinase 1 |
| ACTN1 | Actinin Alpha 1 |
| PAK3 | P21 (RAC1) Activated Kinase 3 |
| ITGB1 | Integrin beta 1 |
| ITGA5 | Integrin alpha 5 |
| Crk | CRK Proto-Oncogene, Adaptor Protein |
| Crkl | CRK like Proto-Oncogene, Adaptor Protein |
| Mapk1 | Mitogen-Activated Protein Kinase 1 |
| Mapk8 | Mitogen-Activated Protein Kinase 8 |
| Pak2 | P21 (RAC1) Activated Kinase 2 |
| Src | SRC Proto-Oncogene, Non-Receptor Tyrosine Kinase |
| Diaph1 | Diaphanous Related Formin 1 |
| Pikfyve | Phosphoinositide Kinase, FYVE-Type Zinc Finger Containing |
| Mylk | Myosin light chain kinase |
| Limk | Lim domain kinase 1 |
| Cfl2 | Cofilin2 |

**Supplemental Figure 6: Comparison of cytoskeletal effectors to validate *in vitro***

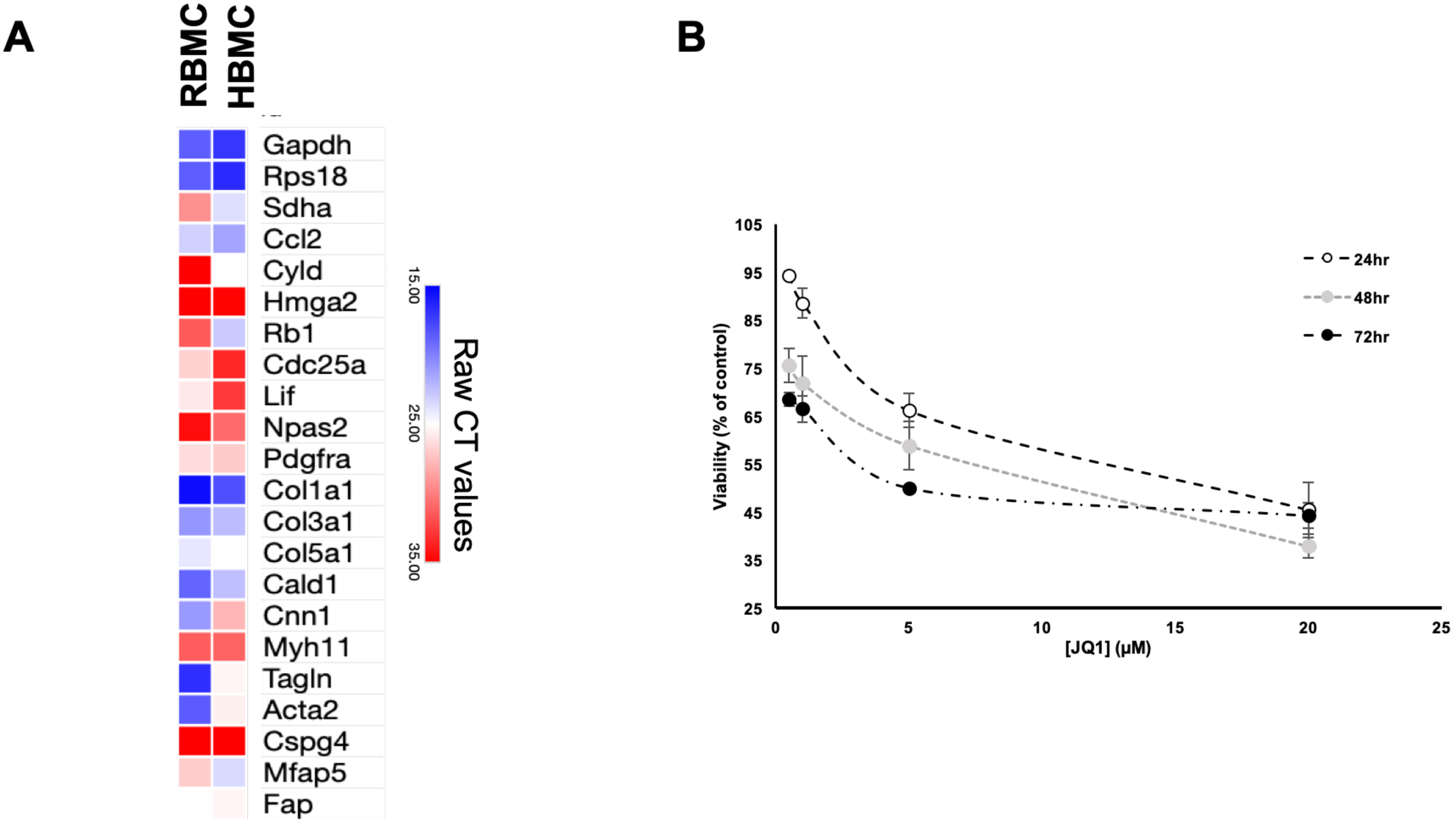


**Supplemental Figure 7: HBSMC and RBMC comparison and JQ1 dose and time effect on cellular viability**

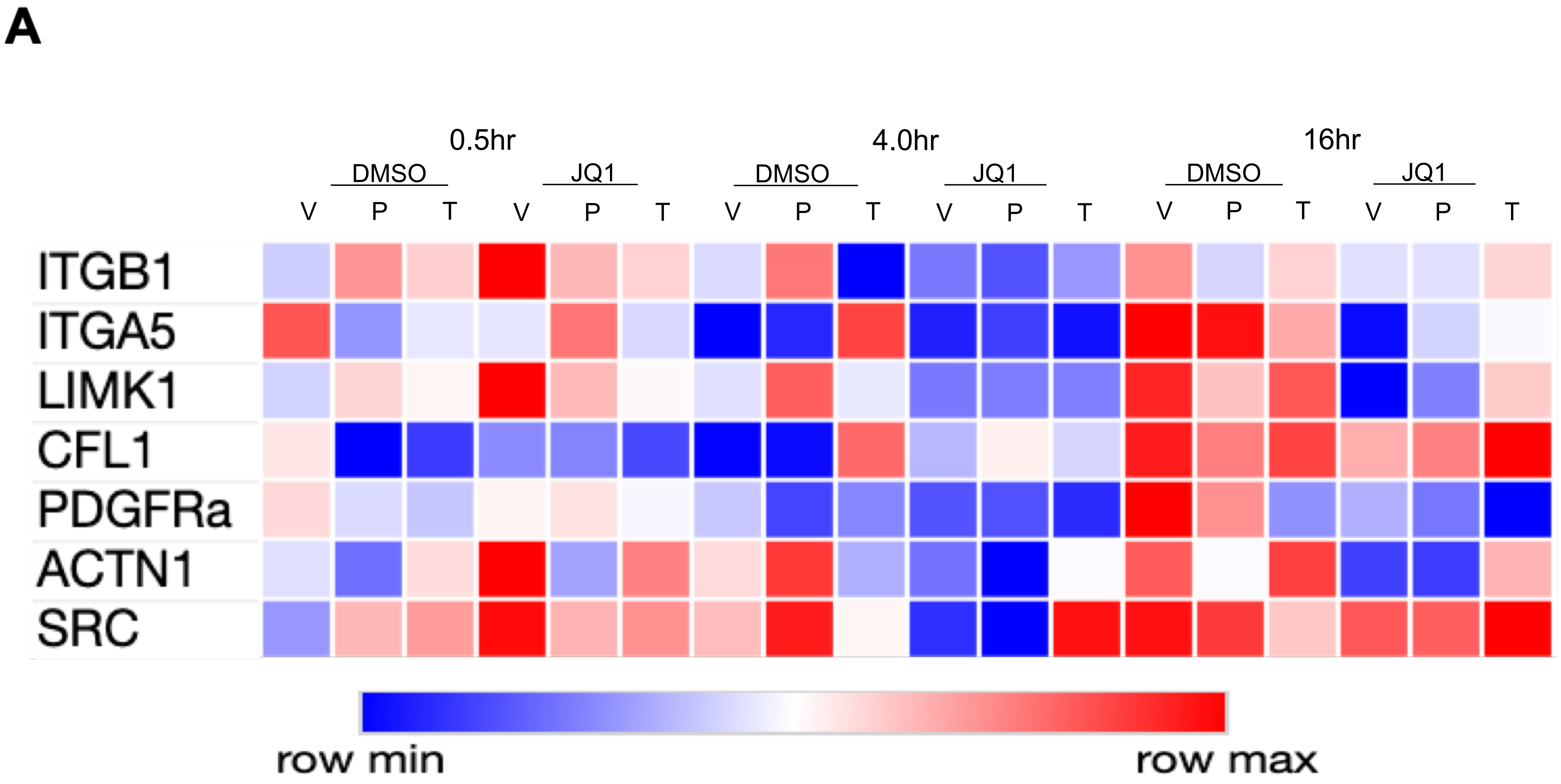


**Supplemental Figure 8: qPCR validation of JQ1-sensitive, cytoskeleton-associated gene in pHBSMC**

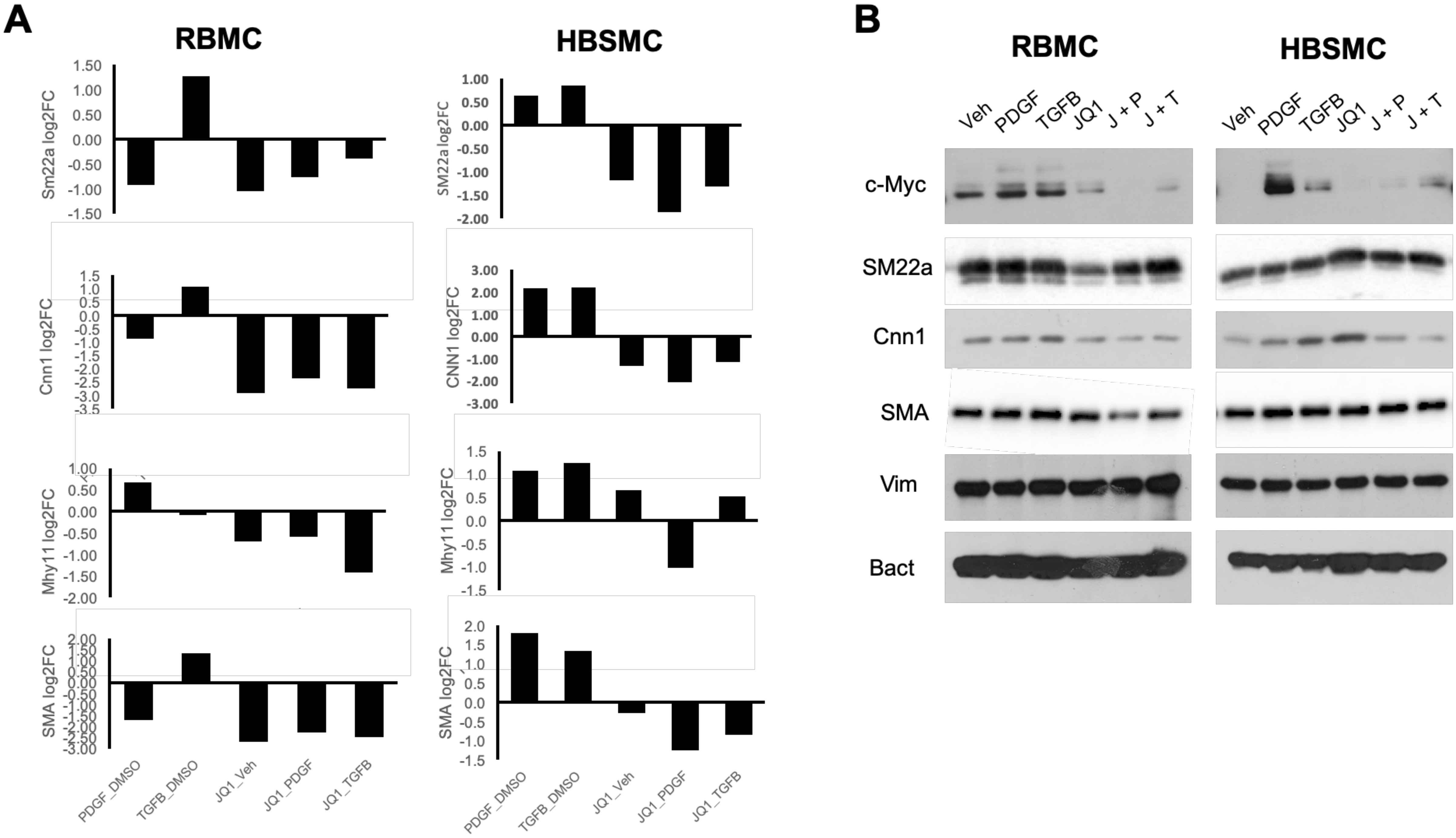


**Supplemental figure 9: JQ1 robustly reduces transcription of contractile genes human and rat bladder contractile cells**

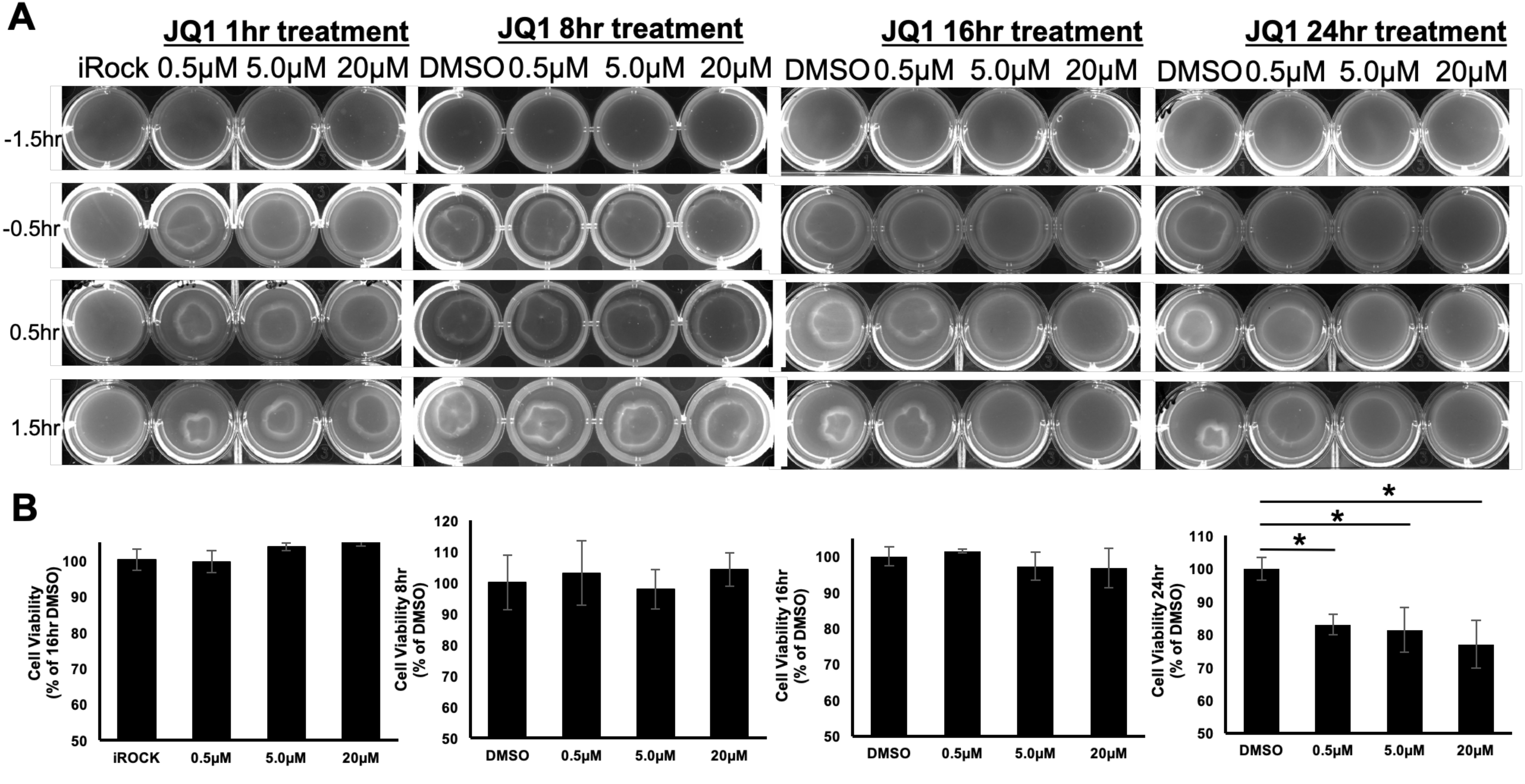


**Supplemental figure 10: JQ1 inhibits contraction of RBMCs on collagen gels after 16hrs with low cytotoxicity**
